# MHCII^+^ cancer-associated fibroblasts promote antitumor immunity in pancreatic cancer

**DOI:** 10.64898/2026.11.03.764075

**Authors:** Jessica Hung, Stephanie Silveria-Davies, Jenna Vickery, Brenna Gittins Maker, Chloe Mikolajczyk, Dorothée Saddier Axe, Brian M. Weist, Joshua E. Elias, Michel DuPage

## Abstract

The functions of major histocompatibility complex class II molecule-expressing cancer-associated fibroblasts (MHCII^+^CAFs) in pancreatic cancer remain controversial. MHCII^+^CAFs are primarily described as immunosuppressive, as MHCII allows for direct interaction with, and support of, regulatory T cells (Tregs). However, MHCII^+^CAF functions may be more complex depending on their origins and their cellular interactions within the tumor microenvironment. Here, using orthotopic models of pancreatic cancer, we show that conventional CD4^+^ T cells and their production of Interferon gamma (IFNγ) increased MHCII^+^CAFs in tumors. Acute depletion of Tregs using *Foxp3^DTR^* mice revealed that Tregs suppress the abundance of MHCII^+^CAFs in tumors by inhibiting CD4 T cell responses that can directly control primary tumor growth and metastatic progression. Functionally, MHCII expression on fibroblasts was required for CD4 T cell control of PDAC progression upon Treg ablation. Collectively, our findings identify an IFNγ-driven, antitumor activity of MHCII expression on CAFs and reveal that CAFs may be harnessed to promote adaptive immunity to PDAC.

## INTRODUCTION

Pancreatic ductal adenocarcinoma (PDAC) is one of the deadliest cancers, with a 5-year survival rate of 13% in 2026^1^. The aggressive nature of PDAC is driven, in part, by its characteristically desmoplastic reaction, exemplified by the recruitment and expansion of a heterogeneous pool of cancer-associated fibroblasts (CAFs) that can directly support cancer cell proliferation and invasion^2–6^. The presence of CAFs in tumors also thwarts many treatment regimens by impeding drug delivery or immune infiltration^7–9^. Complicating our understanding of CAFs in PDAC is their phenotypic heterogeneity, which is clearly visualized by single-cell RNA sequencing studies that have identified distinct CAF populations in murine and human PDAC. Three major CAF populations are widely recognized: myofibroblastic CAFs (myCAFs), inflammatory CAFs (iCAFs), and MHC class II-expressing CAFs (MHCII^+^CAFs, commonly termed antigen-presenting CAFs, apCAFs)^10–15^. Each CAF population exhibits distinct transcriptional programs that may promote or restrain tumor growth. Understanding their subset-specific functions and the signals that drive their abundance in tumors may inform strategies to reprogram CAFs to improve therapeutic outcomes.

In PDAC, the recruitment of immunosuppressive myeloid cells and the accumulation of regulatory T cells (Tregs) can enforce local immune suppression in tumors, promoting tumor progression^16–19^. Notably, increased intratumoral Treg abundance has been associated with adverse clinical outcomes in PDAC^20–22^, and targeting intratumoral Tregs with TNFR2 blockade can enhance antitumor responses in preclinical models^23^. Most recently, several studies have hypothesized that MHCII^+^CAFs in PDAC present antigens on MHCII molecules to CD4^+^ T cells, which selectively supports the differentiation of immunosuppressive Tregs^10,24,25^. However, the capacity of MHCII^+^CAFs to promote Treg differentiation has been performed in vitro, and studies in vivo that test whether MHCII^+^CAFs promote or restrain tumor progression are lacking. In PDAC, MHCII^+^CAFs were first described to arise from a mesothelial lineage, separate from myCAFs and iCAFs^24^. These apCAFs were shown to induce Foxp3^+^Treg differentiation in vitro^24,25^. However, other studies have shown that MHCII expression on CAFs may also be induced, thus the activity of MHCII on CAFs may depend on the origins of the CAF population^14,26,27^. For example, MHCII^+^CAFs in lung cancer promoted

Foxp1^+^Treg expansion through PD-L2 upregulation and impeded the efficacy of chemoimmunotherapy^26^. Conversely, increased MHCII^+^CAF abundance has been associated with favorable prognosis and immunotherapy response in gastric and breast cancers^28^, and loss of MHCII^+^CAFs in pancreatic and lung cancers impaired responsiveness to immune checkpoint blockade immunotherapy^25,27^. Together, these findings highlight context-dependent roles for MHCII^+^CAFs in tumor immunity and therapeutic responses. However, how the immune microenvironment regulates MHCII^+^CAF abundance during tumor progression remains underexplored. Defining the immune cell populations and interactions that govern this fibroblast state in vivo will help explain how immune perturbations reshape CAF composition and influence tumor control.

Here, we investigated how distinct CD4^+^ T-cell populations regulate MHCII^+^CAFs and whether MHCII expression in fibroblasts influences tumor control in vivo. Using orthotopic PDAC models containing high or low levels of T cell infiltration, we found that conventional CD4^+^ T cells (Tconv), but not CD8+ T cells, were required to maintain MHCII^+^CAF abundance in primary tumors. Treg ablation promoted tumor control and also increased MHCII^+^CAF abundance in an IFNγ-dependent manner. Depletion of CD4^+^ Tconv or neutralization of IFNγ abrogated both tumor control and MHCII^+^CAF reprogramming following Treg ablation, suggesting that MHCII expression on CAFs may contribute to tumor control. Indeed, CAF-specific MHCII deficiency diminished tumor control, whereas co-implantation of cancer cells with MHCII^+^ fibroblasts, but not MHCII-deficient fibroblasts, promoted tumor control. These findings were recapitulated in a therapeutically translatable setting of combined anti-PD-1 checkpoint blockade and anti-CCR8-directed intratumoral Treg depletion. Collectively, our findings identify conventional CD4^+^ T cells and IFNγ as positive regulators of MHCII^+^CAFs in PDAC and uncover MHCII expression in CAFs as a promoter of antitumor immunity.

## RESULTS

### Different pancreatic cancer-derived lines generate T cell low or T cell high tumor environments upon orthotopic growth

Previously, a series of distinct tumor lines derived from KPC (*Kras^G12D/+^;Trp53^R172H/+^;Pdx1-Cre*) pancreatic ductal adenocarcinomas (PDAC) in mice were shown to exhibit a wide range of T cell infiltration when subcutaneously transplanted^29^. To test whether the cellular composition of these PDAC clones is conserved when grown within their native tumor microenvironment, we performed orthotopic implantation of four different KPC-derived PDAC lines into the pancreas of syngeneic immunocompetent C57BL/6 mice (**Figure 1A**)^29,30^. Analysis of these tumors sixteen days after orthotopic growth revealed a diversity of immune cell infiltrates by flow cytometry (**Figure 1B and S1A**). Interestingly, tumors most heavily populated by activated T cells correlated with reduced primary tumor size and fewer metastases, although the smaller size of tumors may also be due to an intrinsically slower proliferation rate (**Figure 1C and S1B-C**). Further analyses focused on the PDAC lines 6421c2 and FC1245, which had the strongest and weakest T cell responses in tumors, respectively (**Figure 1D-1F**). Orthotopic 6421c2 tumors, compared to FC1245, also exhibited increased infiltration of dendritic cells (DCs) and NK cells, but reduced regulatory T cells (Tregs) and potentially suppressive myeloid cell populations (neutrophils, Ly6C-hi-, and Ly6c-low-monocytes) (**Figure 1G, 1H, and S1D-G**). These findings reveal distinct immune compositions of PDAC despite being driven by the same underlying genetic events.

**Figure 1.**
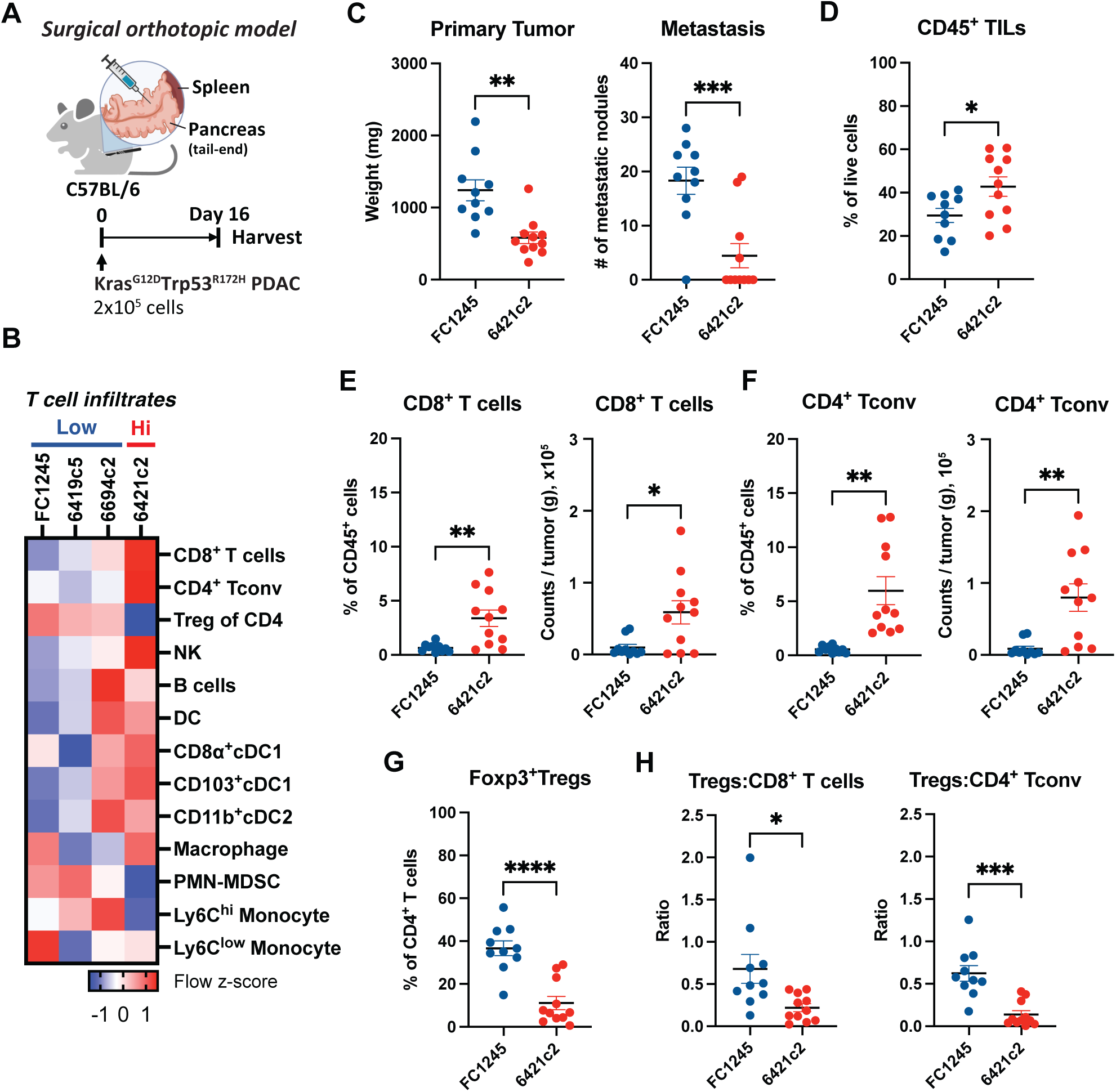
Different pancreatic cancer-derived lines generate T cell-low or T cell-high tumor microenvironments in the orthotopic model. (A) Schematics of the surgical orthotopic PDAC model. KP cell lines were surgically implanted at the tail-end of the pancreas. Tumors were harvested on day 16 post-implantation. (B) Flow cytometric immunophenotyping of tumors derived from four KP PDAC lines at harvest. The heatmap shows z-scores of immune cell population frequencies among CD45^+^ cells. (C) Primary tumor mass and number of metastatic nodules in mice bearing FC1245 or 6421c2 tumors. (D) Frequency of CD45^+^ tumor-infiltrating leukocytes among live cells in FC1245 and 6421c2 tumors. Frequency and numbers per gram of tumor of (E) CD8^+^ T cells and (F) conventional CD4^+^ T cells in FC1245 and 6421c2 tumors. (G) Frequency of Tregs in CD4^+^ T cells in FC1245 and 6421c2 tumors. (H) Ratios of Tregs to CD8^+^ T cells and to conventional CD4^+^ T cells in FC1245 and 6421c2 tumors. (B), Data shown are representative of two independent experiments with the same results. (C-H), Data are pooled from two independent experiments. Data are shown as mean ± SEM, statistical analysis by unpaired Student’s t-test; *p<0.05; **p<0.01; ***p<0.001; ***p<0.0001.

### CAF composition differs in high-and low-T cell-infiltrated tumors

The presence of cancer-associated fibroblasts (CAFs) is associated with tumor progression in PDAC, in part because CAFs can block anti-cancer immune responses^7,31–33^. To test whether differences in CAF composition correlated with immunologically distinct PDAC tumors, we examined the frequencies of the major CAF subsets: myofibroblast CAFs (myCAFs), inflammatory CAFs (iCAFs), and MHC class II-positive CAFs (MHCII^+^CAFs), in high-and low-T cell-infiltrated PDAC tumors by flow cytometry (**Figure 2A**). T cell-high 6421c2 tumors contained the highest frequency of total (podoplanin-positive, PDPN^+^) CAFs compared to all other tumor lines, especially FC1245 (**Figure S2A-C**). Among CAF subsets, all the T cell-low tumors had significantly lower frequencies of MHCII^+^CAFs and iCAFs compared to 6421c2 tumors and were instead significantly enriched in myCAFs (**Figure 2B and S2D**). To determine whether the increase in Ly6C^-^MHCII^-^myCAFs determined by flow cytometry analysis was reflected in situ in tumor tissues, we performed immunofluorescent staining for the myCAF marker alpha-smooth muscle actin (αSMA) and Mason trichrome staining for collagen deposition in tumor sections. Consistent with the flow cytometry data, FC1245 PDAC exhibited a greater αSMA^+^ area and increased collagen compared to 6421c2 PDAC, indicating increased myCAF abundance in T cell-low tumors (**Figure 2C and 2D**). Thus, these data may suggest that increased myCAF abundance in PDAC lines promotes tumor growth and prevents immune cell recruitment, as previously reported^2,3,8,33^. However, these results may also implicate MHCII^+^CAFs or iCAFs in the promotion of T cell responses and tumor control in the 6421c2 PDAC line.

**Figure 2.**
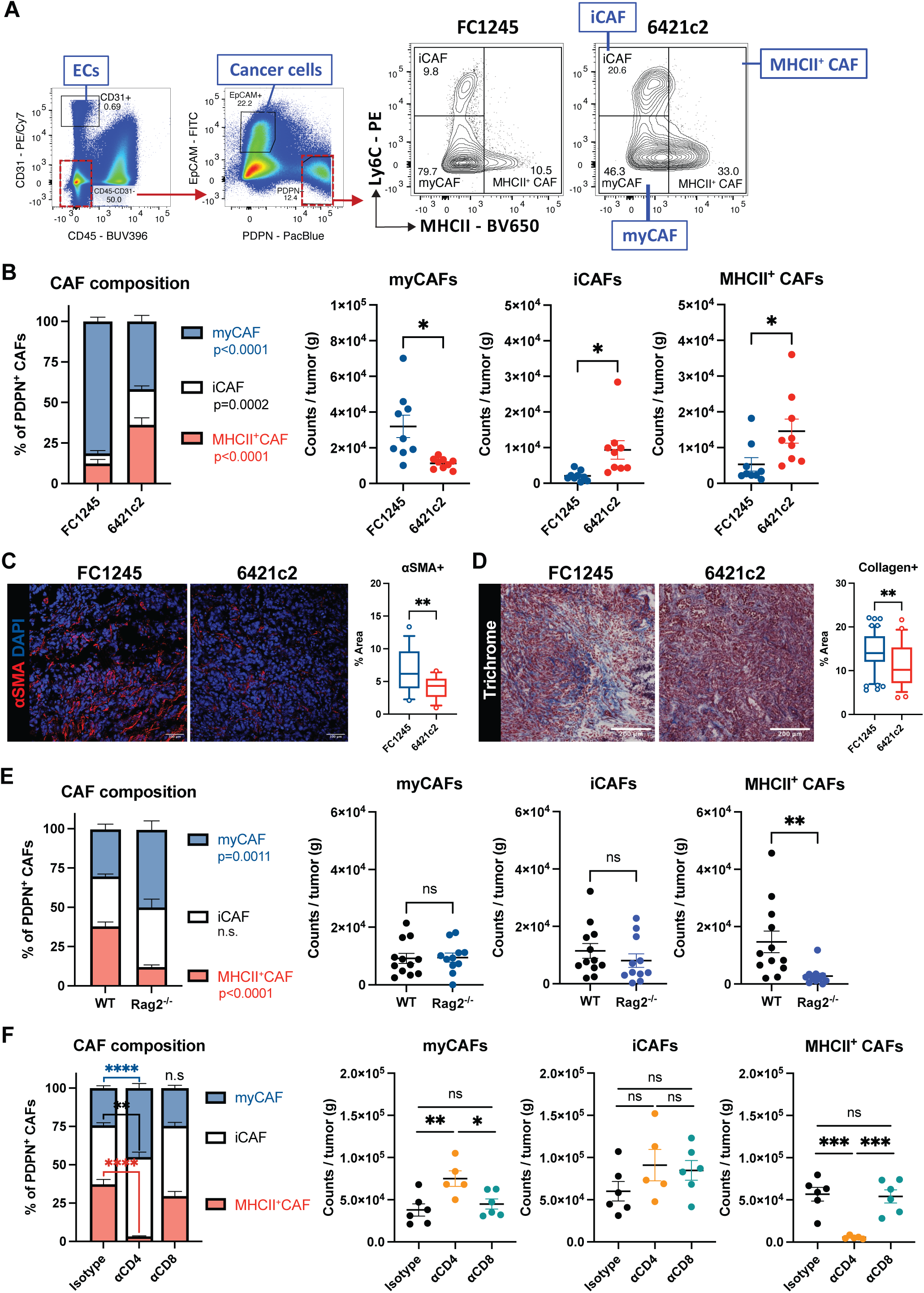
The abundance of MHCII^+^CAF is dependent on intratumoral CD4^+^ T cells. (A) Gating strategy for three CAF subsets, myCAFs (Ly6C^-^MHCII^-^), iCAFs (Ly6C^+^MHCII^-^), and MHCII^+^CAFs (MHCII^+^). (B) Frequencies of CAF subsets among PDPN^+^CAFs and numbers per gram of tumors in FC1245 and 6421c2 tumors on day 16 post-implantation. (C) Representative immunofluorescent image and quantification of αSMA staining in FC1245 and 6421c2 tumors. Red, αSMA; Blue, DAPI; scale bar, 100 μm. Boxplot shows the 10^th^-90^th^ percentiles. (D) Representative Mason trichrome staining of FC1245 and 6421c2 tumors. Scale bar, 200 μm. Boxplot shows the 10^th^-90^th^ percentiles. (E) CAF subset composition of orthotopic 6421c2 tumors in wild-type and *Rag2^-/-^*mice on day 16 post-implantation. (F) CAF subset composition of orthotopic 6421c2 tumors treated with IgG2b, anti-CD4, or anti-CD8 antibodies on day 16 post-implantation. Data are pooled from two independent experiments. Data are shown as mean ± SEM; *p<0.05; **p<0.01; ***p<0.001; ***p<0.0001. (B, E), Frequency: Ordinary Two-way ANOVA with Šídák’s multiple comparison test; number per gram tumor: unpaired Student’s t-test. (C, D), unpaired Student’s t-test (n=4-8 slides/mouse, 5 mice/group). (F), Frequency, Ordinary Two-way ANOVA with Šídák’s multiple comparison test; number per gram tumor, One-way ANOVA with Tukey’s multiple comparison test.

### The presence of MHCII^+^CAFs is dependent on intratumoral CD4^+^ T cells

To determine whether CAF composition instructs the immune cell populations in PDAC or whether T cell infiltration of PDAC regulates CAF composition, we compared CAF subsets in orthotopic 6421c2 tumors inoculated in wild-type mice versus *Rag2^-/-^* immunodeficient mice, which lack T and B cells. In tumors from *Rag2^-/-^* mice, MHCII^+^CAFs were significantly reduced compared to tumors from WT mice, while the numbers of myCAFs and iCAFs were unchanged in the presence or absence of T and B cells (**Figure 2E and S2E**). This suggests that cells of the adaptive immune system may generate or maintain MHCII^+^CAFs in PDAC.

Next, using antibody depletion of either CD4^+^ or CD8^+^ T cell subsets in 6421c2 tumor-bearing mice, we showed that only CD4^+^ T cell depletion recapitulated the results observed in *Rag2^-/-^* mice, significantly reducing MHCII^+^CAFs in tumors (**Figure 2F, S2F and S2G**). These findings suggest that CD4^+^ T cells promote MHCII^+^CAF abundance in PDAC. It is noteworthy that changes in CAF composition and the presence or absence of T cells did not impact the progression of 6421c2 (**Figure S2E and S2H**). However, this is in line with previous observations made with these PDAC lines in the setting of subcutaneous tumor growth, where T cells only controlled tumor progression with the addition of immunotherapeutic approaches.

### Ablation of regulatory T cells promotes tumor control and the upregulation of MHCII^+^CAFs

To distinguish the contribution of conventional CD4^+^ T cells versus suppressive Tregs in altering the CAF composition in PDAC, especially because MHCII^+^CAFs have been reported to interact with Tregs in PDAC^24,25^, we used *Foxp3^DTR^*mice to specifically ablate Tregs and assess changes to CAF populations in PDAC. First, we confirmed that systemic administration of diphtheria toxin (DT) in established orthotopic PDAC achieved complete intratumoral Treg ablation in both FC1245 and 6421c2 tumors (**Figure 3A, S3A, and S3B**). Second, total Treg ablation in this short-term setting occurred without inducing significant signs of autoimmunity – 75% of DT-treated mice (8/10 for FC1245 and 7/10 for 6421c2) maintained >85% of their initial body weight throughout the treatment course (**Figure S3C**). Finally, as Treg ablation is an effective immunotherapeutic cancer treatment in multiple mouse cancer models^34–38^, DT treatment in this setting also significantly controlled primary 6421c2 and FC1245 PDAC, as well as reduced metastases from FC1245 (**Figure 3B**). Ultimately, Treg ablation led to significantly prolonged survival in both 6421c2- and FC1245-bearing mice (**Figure 3C**).

**Figure 3.**
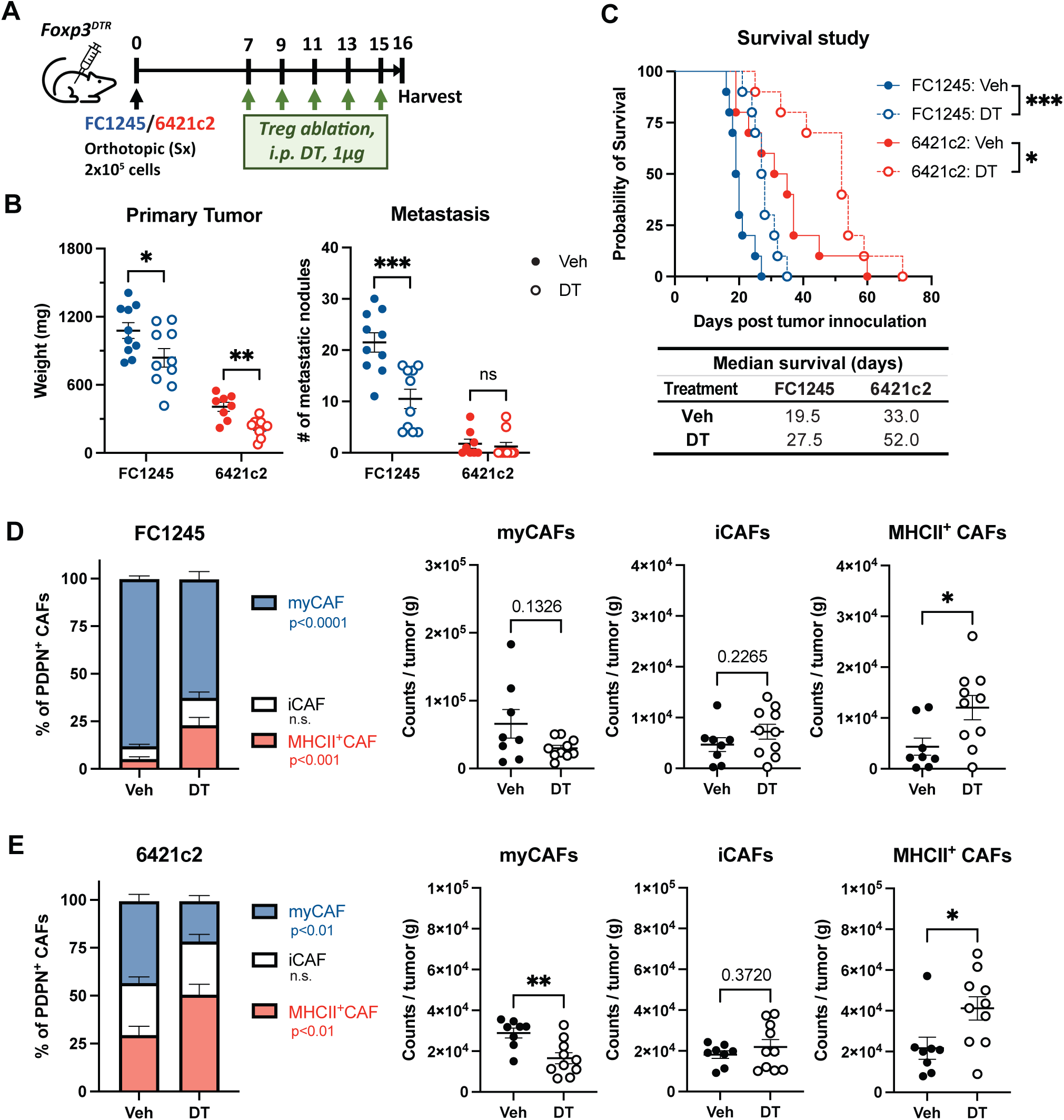
Treg ablation promotes tumor control and upregulation of MHCII^+^CAFs. (A) Schematic of Treg ablation in *Foxp3^DTR^* mice with surgical (sx) implantation of FC1245 or 6421c2 in the pancreas. (B) Primary tumor mass and number of metastatic nodules in FC1245 and 6421c2 tumors following Treg ablation. (C) Survival study of FC1245-and 6421c2-bearing mice receiving continuous DT injections starting on day 7 post-tumor implantation. CAF subset composition in (D) FC1245 and (E) 6421c2 tumors in Treg-ablated mice. Data are pooled from two independent experiments. Data are shown as mean ± SEM; *p<0.05; **p<0.01; ***p<0.001. (B), unpaired Student’s t-test comparing vehicle vs DT within the same tumor model. (C), Mantel-Cox Logrank test (n=10 mice/group). (D, E), Frequency, Ordinary Two-way ANOVA with Šídák’s multiple comparison test; number per gram tumor, unpaired Student’s t-test.

With respect to CAF populations, Treg ablation in both PDAC models led to a significant increase in both the frequency and number of MHCII^+^CAFs in tumors (**Figure 3D, 3E, and S3D**). However, Ki67 staining was not increased in all CAF populations with Treg ablation (**Figure S3E-G**). This suggests that Treg ablation may lead to the reprogramming of resident MHCII-negative fibroblasts to express MHCII rather than cause the selective expansion of pre-existing MHCII^+^CAFs. In contrast to increased MHCII^+^CAFs, we observed a significant reduction of myCAFs in DT-treated 6421c2 and a trend towards fewer myCAFs in FC1245 tumors. These reductions in myCAFs with Treg ablation are supported by studies that identified Tregs as a critical source of TGF-β that supports myCAF maintenance in PDAC^17,39^. However, our results with Treg ablation (**Figure 3E**) and total CD4 T cell depletion (**Figure 2F**) might also suggest that the activity of Tconv cells could negatively regulate the abundance of myCAFs in tumors. Finally, no significant changes in iCAF abundance were observed following Treg ablation. Thus, the presence of Tregs in PDAC appears to prevent the accumulation or maintenance of MHCII^+^CAFs and suggests that CD4^+^ Tconv cells may act to promote MHCII^+^CAFs, as well as reduce myCAFs, in PDAC.

### The presence of MHCII^+^CAFs is associated with the success of antibody-mediated cancer immunotherapies

Next, we tested clinically translatable immunotherapeutic approaches to deplete intratumoral Tregs (anti-CCR8) alone or in combination with checkpoint blockade (anti-PD1)^40–42^. Anti-CCR8 treatment induced an approximate 70% reduction in intratumoral Tregs in 6421c2 PDAC, with no significant changes in tumor-draining and non-draining LNs (**Figure S4A-C**). While intratumoral Treg ablation with anti-CCR8 was not as complete as with DT treatment in *Foxp3^DTR^* mice, unlike the *Foxp3^DTR^* model (**Figure S3A and S3B**), anti-CCR8 treatment alone or in combination with anti-PD-1 exhibited less toxicity, as anti-CCR8-treated mice did not lose any weight (**Figure S4D**). With respect to tumor control, while anti-CCR8 or anti-PD-1 treatments alone produced a similar reduction in the size of primary tumors (**Figure S4E**), the combination of anti-CCR8 and anti-PD-1 achieved the most robust control of primary PDAC (**Figure S4E**). Interestingly, the combination of anti-CCR8 and anti-PD-1 also significantly increased MHCII^+^CAFs and reduced myCAFs in tumors (**Figure S4F**). Taken together, these findings reveal a strong correlation between the presence of MHCII^+^CAFs and improved immunotherapeutic control of orthotopic PDAC.

### Tregs suppress conventional CD4^+^ T cell-dependent induction of MHCII^+^CAFs and tumor control

The opposing effects of total CD4^+^ T cell and Treg depletion on MHCII^+^CAF frequency led us to hypothesize that Tregs regulate MHCII^+^CAFs indirectly, most likely by suppressing CD4^+^ Tconv cells. Indeed, following DT treatments, we observed increased frequencies of both CD4^+^ Tconv and CD8^+^ T cells within tumors (**Figure S3A and S3B**). To test whether the increase in MHCII^+^CAFs upon Treg ablation was due to the activity of CD4^+^ or CD8^+^ T cells, we again used antibody-mediated cellular depletions (**Figure 4A**). Interestingly, CD4, but not CD8, T cell depletion blocked the increase in MHCII^+^CAFs following Treg ablation, suggesting that Tregs do suppress MHCII^+^CAFs indirectly by inhibiting the activity of CD4^+^ Tconv cells (**Figure 4B-D**). Consistent with this finding, geometric mean fluorescence intensity (gMFI) analysis demonstrated that MHCII expression on PDPN^+^CAFs was dependent on CD4^+^ Tconv cell abundance, irrespective of cancer cell-intrinsic differences between two PDAC clones (**Figure 4E**). Like MHCII^+^CAF abundance, tumor control was abrogated in both FC1245 and 6421c2 models only by depleting CD4^+^ Tconv cells (**Figure 4F and 4G**). The importance of CD4 T cell-dependent tumor control with Treg ablation is consistent with our previous findings that CD4^+^ Tconv cells, but not CD8^+^ T cells, were required for the control of MHC I-sufficient and -deficient subcutaneous MC38 and B16F10 tumors^34,35^. Together, these findings demonstrate that CD4^+^ Tconv cells are required for MHCII^+^CAF reprogramming and tumor control following Treg ablation in PDAC. These results may also imply an important role for MHCII^+^CAFs in supporting immunotherapeutic tumor control.

**Figure 4.**
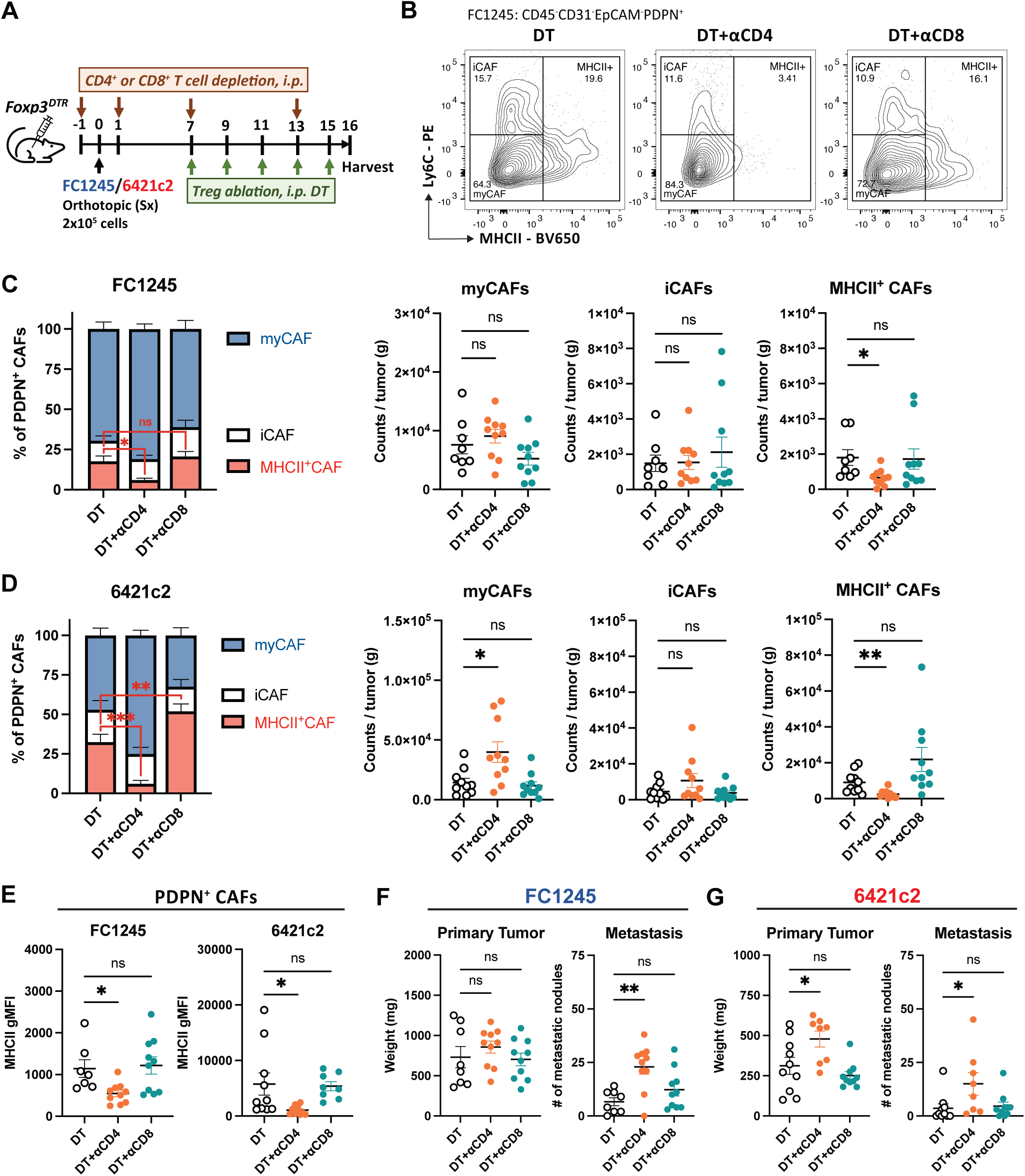
Tregs suppress conventional CD4^+^ T cell-dependent tumor control and induction of MHCII^+^CAFs. (A) Schematic of CD4-or CD8-depletion in Treg-ablated, FC1245- and 6412c2-bearing *Foxp3^DTR^* mice. (B) Representative flow plots of CAF subsets in FC1245 tumors treated with DT plus IgG2b, anti-CD4, or anti-CD8 antibodies. Frequencies and number of CAF subsets per gram of (C) FC1245 and (D) 6421c2 tumors with T cell depletion in Treg-ablated models. (E) MHCII expression in gMFI of PDPN^+^CAFs in FC1245 and 6421c2 tumors. Primary tumor mass and number of metastatic nodules in the (F) FC1245 and (G) 6421c2 models following treatment with DT plus IgG2b, anti-CD4, or anti-CD8 antibodies. Data are pooled from two independent experiments. Data are shown as mean ± SEM; *p<0.05; **p<0.01. (B, C, F, G, H), One-way ANOVA with Tukey’s multiple comparison test. (E, G), Ordinary Two-way ANOVA with Šídák’s multiple comparison test.

### IFNγ-secreting Th1s are required for MHCII^+^CAF reprogramming and tumor control

Next, to investigate how CD4^+^ Tconv cells promote MHCII^+^CAF reprogramming and tumor control, we measured cytokine concentrations from single-cell suspensions of PDAC tumors. Cytokines are known to be important in CD4-dependent tumor control and have been shown to drive the differentiation of CAF states^9,43^. A flow-based multiplex ELISA measuring 13 common proinflammatory cytokines identified IFNγ as uniquely and significantly increased in T cell high and low PDAC models with Treg ablation (**Figure 5A and S5A**). Importantly, increased IFNγ was prevented by the depletion of CD4^+^ Tconv cells, which may suggest that CD4^+^ Tconv cells are the source of IFNγ. In contrast, no significant differences were observed in the amount of IFNβ, TNFα, IL-1α, IL-1β, IL-6, IL-10, IL-12p70, IL-17A, IL-23, GM-CSF, and MCP-1 following Treg or Treg plus CD4^+^ T cell depletion (**Figure S5B-L**). Direct intracellular cytokine staining for IFNγ confirmed that CD4^+^ Tconv cells produced more IFNγ upon Treg ablation (**Figure 5B and 5C**). The CD4^+^ Tconv-dependent increase in IFNγ suggested Tregs may suppress a specific CD4 differentiation program. Indeed, we observed a significant increase in T-bet^+^ Th1 cells among activated (CD62L^-^CD44^+^) CD4^+^ Tconv cells in DT-treated 6421c2 tumors, as well as in tumor-draining and non-draining lymph nodes (**Figure 5D and 5E**). To a lesser extent, GATA3^+^ Th2 cells were also increased in DT-treated mice, while no differences were observed in RORγt^+^ Th17 or in CD4^+^ Tconv cells expressing NK cell receptors NKG2A/C/E^44,45^. Consistent with the findings in 6421c2 tumors, Th1 cells also trended towards increased abundance in DT-treated FC1245 tumors (**Figure S6**).

**Figure 5.**
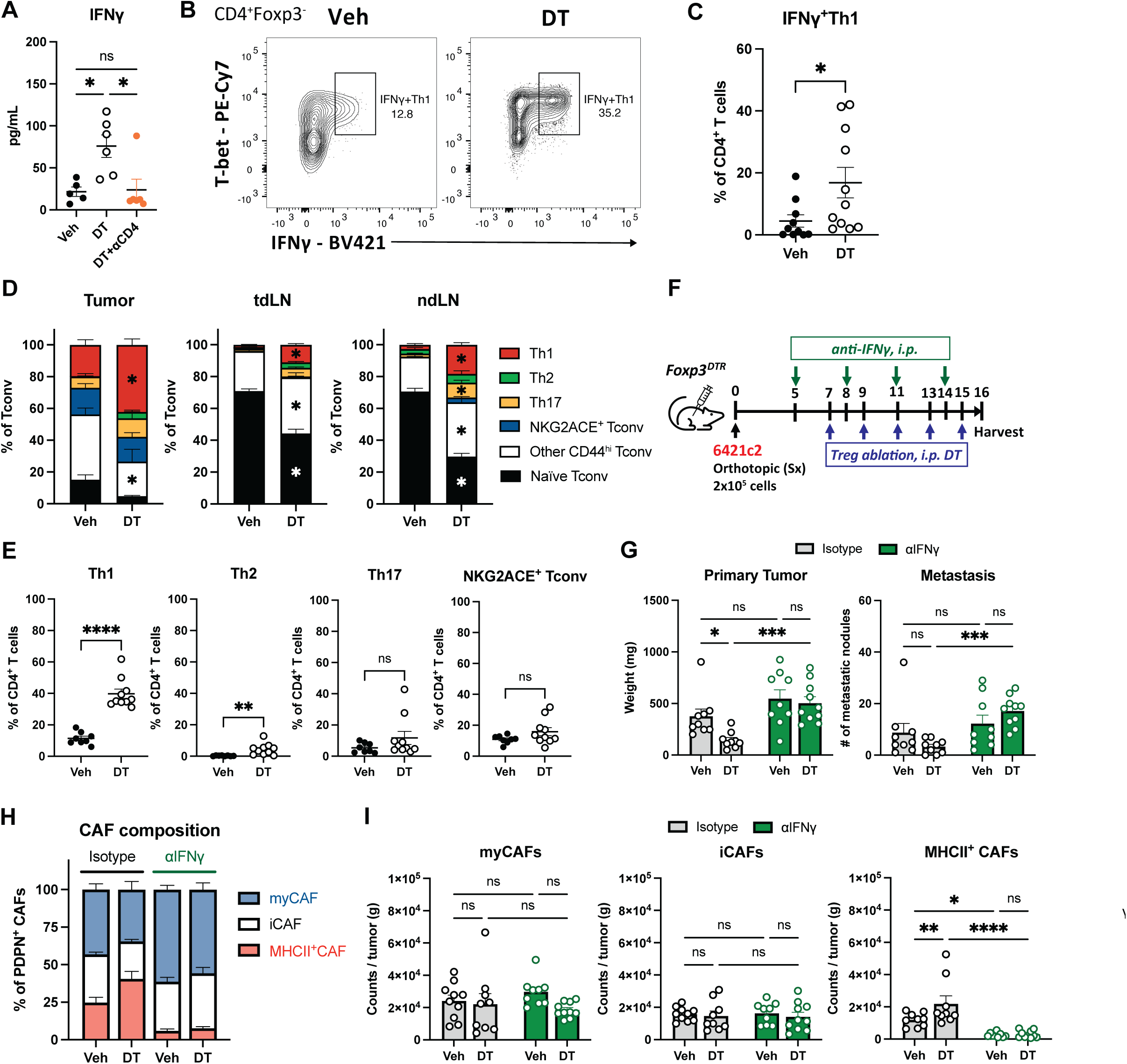
IFNγ-secreting Th1s are required for MHCII^+^CAFs reprogramming and tumor control. (A) IFNγ concentration in 6421c2 tumors treated with vehicle, DT + IgG2b, or DT + anti-CD4 antibodies. (B) Representative flow plots of IFNγ-secreting T-bet^+^Th1 in 6421c2 with or without Treg ablation. (C) Frequency of IFNγ^+^T-bet^+^Tconv among CD4^+^ T cells in 6421c2 +/-DT. (D) Composition of CD4^+^Tconv subsets in tumor, tdLN, and ndLN of 6421c2 +/- DT. (E) Frequency of Th1(T-bet^+^), Th2(GATA3^+^), Th17(RORγt^+^), and NKG2ACE^+^ Tconv in CD4^+^ T cells in 6421c2 +/- DT. (F) Schematics of IgG1 or anti-IFNγ antibody treatments in 6421c2-bearing *Foxp3^DTR^* mice. (G) Primary tumor mass and number of metastatic nodules in 6421c2 model treated with vehicle, DT, vehicle + anti-IFNγ, or DT + anti-IFNγ. CAF subset composition in (H) frequency of PDPN+CAFs and (I) count per gram of tumor in the 6421c2 model treated with vehicle, DT, vehicle + anti-IFNγ, or DT + anti-IFNγ. (B, C), Data shown are representative of two independent experiments with the same results. (C-I), Data are pooled from two independent experiments. Data are shown as mean ± SEM; *p<0.05; **p<0.01; ***p<0.001; ****p<0.0001. (A), One-way ANOVA with Tukey’s multiple comparison test. (C, E), unpaired Student’s t-test. (D, G, H, I), Ordinary Two-way ANOVA with Šídák’s multiple comparison test.

Finally, to test whether IFNγ is required for tumor control and CAF reprogramming, we administered IFNγ-neutralizing antibodies to DT-treated, 6421c2-bearing mice (**Figure 5F**). IFNγ neutralization abrogated the control of both primary and metastatic tumors despite Treg ablation (**Figure 5G**). Additionally, IFNγ neutralization reduced MHCII^+^CAF frequency at baseline and prevented the increase in MHCII^+^CAFs following Treg ablation but had no impact on the abundance of myCAFs or iCAFs (**Figure 5H and 5I**). These findings identify Th1-derived IFNγ as the key regulator of MHCII^+^CAF reprogramming in PDAC, which is also associated with tumor control.

### MHC Class II Expression in CAFs Contributes to Metastatic Control

Although our data demonstrated a positive correlation between MHCII^+^CAF abundance and tumor control, it is unclear whether MHCII^+^CAFs directly contribute to tumor control or instead represent bystanders responding to IFNγ. More importantly, while it is presumed that MHCII expression on CAFs allows for direct interaction with T cell receptors of CD4^+^ T cells, whether MHCII expression on CAFs, per se, is required for tumor control is untested. To directly examine the functional role of MHCII expression in fibroblasts, we generated bone marrow chimeric mice in which MHCII^WT^ or MHCII^KO^ (*I-A^b-/-^*) hosts were reconstituted with MHCII^WT^ bone marrow to restore MHCII expression in professional APCs, e.g., DCs, macrophages, and B cells, while preserving MHCII-deficiency in non-hematopoietic stromal compartments, most importantly, fibroblasts and endothelial cells (ECs) (**Figure 6A, 6B, and S7A**). As CD4^+^ T cells are absent in MHCII^KO^ mice due to impaired positive selection without MHCII expression in the thymic epithelium (**Figure S7B**), we adoptively transferred CD4^+^ T cells from MHCII^WT^ donors after successful bone marrow reconstitution, just prior to orthotopic tumor implantation (day -5). This generated mice with CD4^+^ T cells that repopulated 6421c2 tumors at frequencies comparable to those observed in MHCII^WT^ hosts (**Figure S7C**). In this setting, we observed that tumors with MHCII-deficient CAFs/stroma had significantly greater numbers of metastatic nodules in the liver, lung, and intestines than tumors with MHCII-sufficient CAFs/stroma (**Figure 6C**). While there was no difference in primary tumor size between mice, this may be due to technical limitations of this model to completely reconstruct a CD4^+^ Tconv cell population that can rapidly and robustly restrict primary tumors. These findings suggest that MHCII expression in non-hemopoietic stromal cells, including fibroblasts and endothelial cells, promotes the control of metastases during PDAC progression.

**Figure 6.**
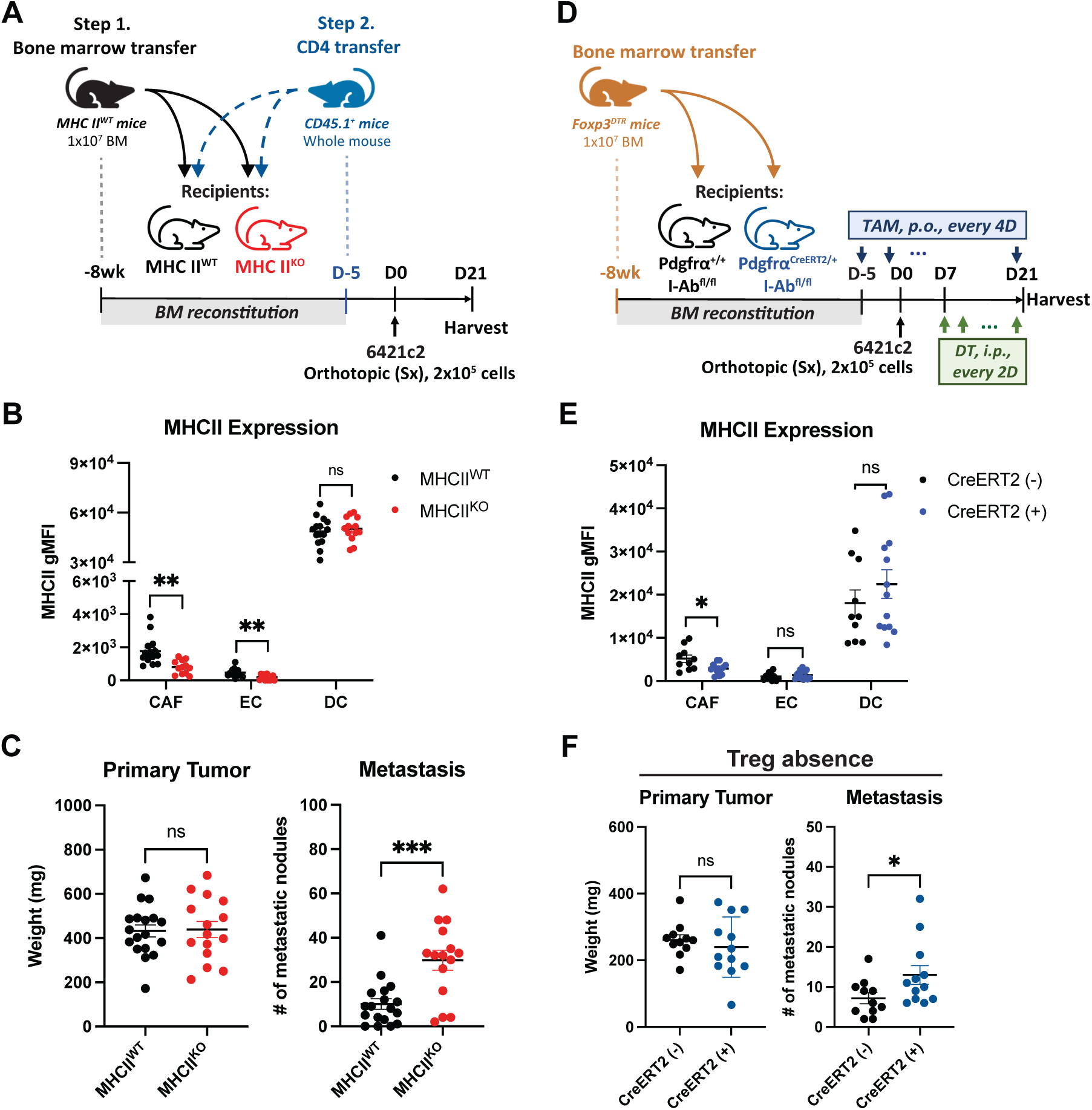
MHC class II expression in CAFs is required for metastatic control. (A) Schematic of the MHCII deficiency model in non-hematopoietic cells in 6421c2-bearing mice. Tumors were harvested on day 21 post-implantation. (B) MHCII gMFI in CAFs, endothelial cells, and dendritic cells in tumors of the stromal MHCII deficiency model at harvest. (C) Primary tumor mass and number of metastatic nodules in 6421c2 tumors with MHCII^WT^ or MHCII^KO^ stroma. (D) Schematic of fibroblast-specific MHCII deletion in 6421c2-bearing mice. Bone marrow from *Foxp3^DTR^* mice was transferred into *Pdgfra^CreERT^*^2^*^/+;^I-Ab^fl/fl^* or *Pdgfra^+/+^;I-Ab^fl/fl^*mice to allow Treg ablation by DT injections. Tumors were harvested on day 21 post-implantation. (E) MHCII gMFI in CAFs, endothelial cells, and dendritic cells in tumors of the fibroblast-specific MHCII deletion model at harvest. (F) Primary tumor mass and number of metastatic nodules in 6421c2 tumors containing MHCII^WT^ (CreERT2-negative) or MHCII^KO^ (CreERT2-positive) CAFs following DT injections. Data are pooled from two independent experiments. Data are shown as mean ± SEM; *p<0.05; **p<0.01; ***p<0.001. (B, E), unpaired Student’s t-test comparing two recipient mice within the same cell type. (C, F), unpaired Student’s t-test.

To directly test whether metastatic control depends on MHCII expression in CAFs, we generated *Pdgfrα^CreERT^*^2^*^/+^;I-Ab^fl/fl^* mice to induce fibroblast-specific deletion of MHCII following Tamoxifen (TAM) administration by oral gavage (**Figure 6D**). Following TAM treatment, CreERT2 positive (+) mice showed a significant reduction in MHCII expression specifically in CAFs compared with CreERT2-negative (-) littermate controls (**Figure 6E**). We next generated bone marrow chimeric mice using bone marrow from *Foxp3^DTR^* mice to enable Treg ablation by DT injections (**Figure 6D**). Consistent with the phenotype observed in mice with MHCII deficiency across all stromal cells (**Figure 6C**), loss of MHCII expression only in CAFs resulted in increased metastatic burden following Treg ablation (**Figure 6F**). Notably, we again did not observe significant differences in primary tumor control, or the control of metastases when Tregs were not ablated (**Figure S7D**). This could reflect incomplete MHCII deletion in the CreERT2-loxP system (**Figure 6E**). However, this result is similar to what we observed previously in mice without Treg ablation (**Figure S2E and S2H**), where the absence of CD4^+^ Tconv cells was associated with fewer MHCII^+^CAFs, but there was no effect on tumor burden. Thus, additional immune perturbation, such as Treg ablation, may be required to reveal the functional contribution of MHCII^+^CAFs to tumor progression.

As noted, MHCII^+^CAFs have been implicated as key drivers of intratumoral immunosuppression in PDAC because in vitro experiments have shown that naïve CD4^+^ T cells stimulated by MHCII^+^CAFs preferentially become Tregs in this setting^24,25^. To test whether MHCII^+^CAFs promoted Treg induction or maintenance within PDAC in vivo, we compared Treg frequencies in tamoxifen-treated *Pdgfrα^+/+^;I-Ab^fl/fl^* versus *Pdgfrα^CreERT^*^2^*^/+^;I-Ab^fl/fl^* mice in the absence of DT to ablate Tregs. Unexpectedly, we observed no difference in intratumoral Treg frequency between tumors with MHCII-sufficient versus MHCII-deficient CAFs, suggesting that MHCII expression on CAFs is not the primary driver of, or is not required for, intratumoral Treg accumulation in vivo (**Figure S7E**). Together, these data demonstrate that the intrinsic expression of MHCII on CAFs is functionally important, and that without it, tumor progression to metastases is increased. Thus, MHCII expression on CAFs may not simply be associated with Treg induction and immunosuppression in PDAC, but may also support effective immunity against PDAC, especially when therapeutically targeting Tregs.

### Addition of MHCII^+^CAFs is sufficient to reduce primary and metastatic tumor growth

The requirement for MHCII expression in CAFs for the control of metastatic PDAC prompted us to ask whether MHCII^+^CAFs are also sufficient to promote tumor control. To test this hypothesis, we generated primary fibroblasts from the lungs of MHCII-deficient and MHCII-sufficient mice^46^. To recapitulate the MHCII^+^CAF state induced in vivo, we pre-stimulated primary fibroblasts with IFNγ (20 ng/mL) in vitro and confirmed induction of MHCII expression in ex vivo-expanded fibroblasts derived from MHCII-sufficient mice (**Figure 7A**). Next, we co-injected IFNγ-stimulated MHCII-deficient or MHCII-sufficient primary fibroblasts with 6421c2 orthotopically at a 5:1 fibroblast-to-cancer cell ratio to evaluate the impact of fibroblasts with or without MHCII on tumor progression (**Figure 7B**). Cancer cells co-implanted with MHCII-negative fibroblasts led to significantly larger tumors and increased metastases compared to mice implanted with cancer cells alone (**Figure 7C**). In contrast, co-implantation with MHCII^+^ fibroblasts significantly reduced both primary and metastatic tumor burden compared with MHCII-negative fibroblast co-implantation, resulting in tumor burdens that were not significantly different from those observed in mice implanted with cancer cells alone (**Figure 7C**). The concurrent recruitment of endogenous CAFs during tumor progression led to a complete replacement of the transplanted CAFs by experimental endpoints, likely preventing any observable changes in the abundance and phenotype of T cells^2^. Nevertheless, these data suggest that transient exposure to MHCII^+^CAFs during the early stages of tumor development is sufficient to promote enhanced tumor control. Overall, these data further support an intrinsic role for MHCII expression in CAFs that can promote, rather than only hinder, the anti-cancer immune response in the setting of PDAC.

**Figure 7.**
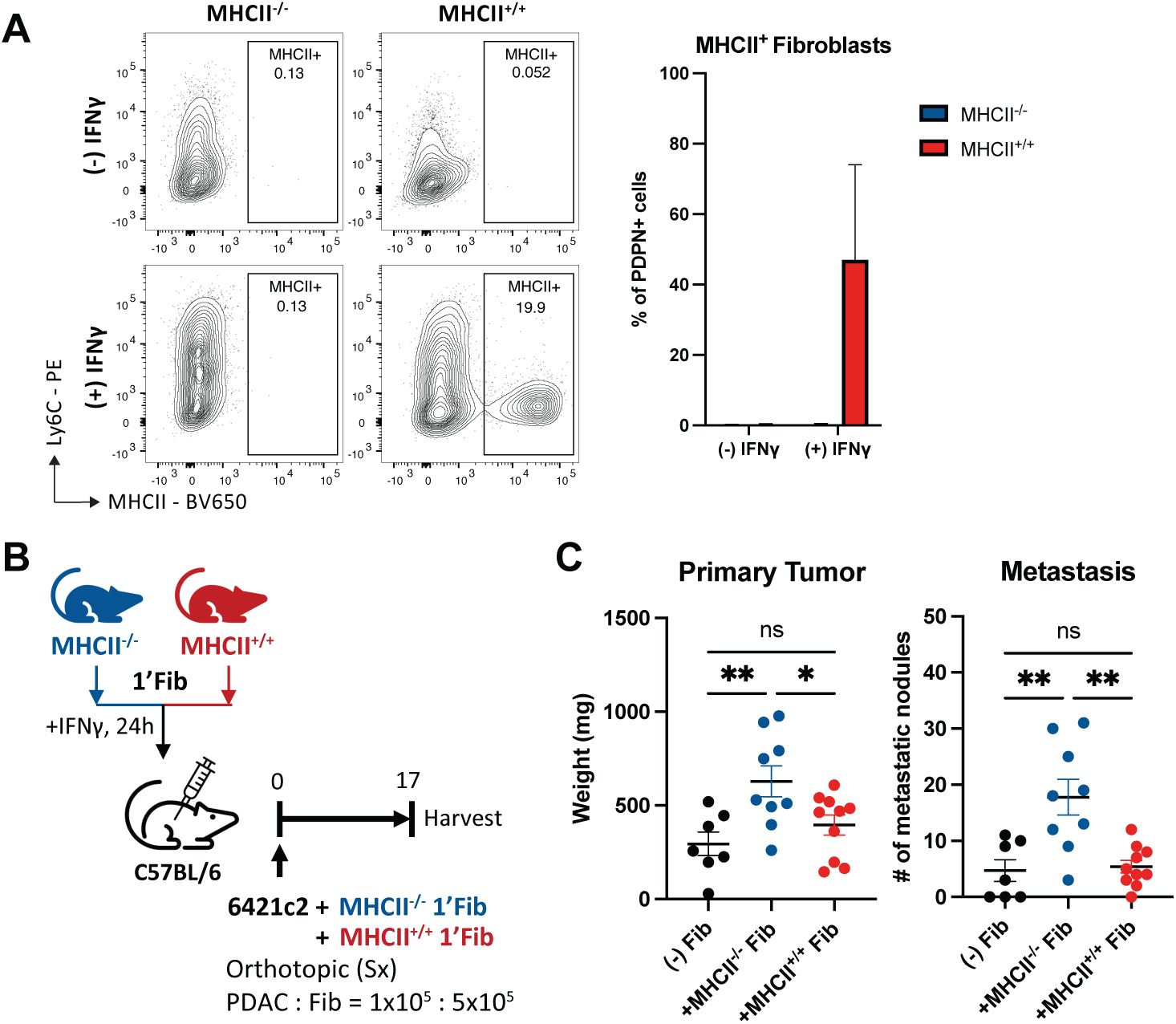
Addition of MHCII^+^CAFs is sufficient to reduce primary and metastatic tumor growth. (A) Representative flow plots and quantification of lung-derived primary fibroblasts from *MHCII^-/-^*or *MHCII^+/+^* mice following IFNγ stimulation, 24h. (B) Schematics of 6421c2 and fibroblast co-implantation models. Tumors were harvested on day 17 post-implantation. (C) Primary tumor mass and number of metastatic nodules in 6421c2 tumors alone, or co-implantation with MHCII-negative fibroblasts or with MHCII-positive fibroblasts. Data are pooled from two independent experiments. Data are shown as mean ± SEM; *p<0.05; **p<0.01. (C), One-way ANOVA with Tukey’s multiple comparison test.

## DISCUSSION

Although previous studies have demonstrated that MHCII⁺CAFs modulate CD4⁺ T cell responses in vitro^10,24,25^, how immune signals regulate fibroblast MHC class II expression and how this expression contributes to tumor progression in vivo remain poorly defined. Using orthotopic PDAC models with Treg ablation and antibody-mediated cellular depletion or cytokine blockade, we identified conventional CD4⁺ Tconv cells and IFNγ as essential regulators of MHCII⁺CAF reprogramming in PDAC. Treg ablation promoted Th1 polarization and increased IFNγ production by conventional CD4⁺ T cells, connecting CD4^+^ T cell responses to stromal remodeling. Consistent with findings in lung cancer^26,27^, IFNγ was required for MHCII^+^CAF reprogramming in pancreatic tumors. Although both activated CD8⁺ and CD4⁺ T cells can produce IFNγ^47^, we found that MHCII⁺CAF reprogramming required conventional CD4⁺ T cells, but not CD8⁺ T cells, across two distinct PDAC clones. Functionally, loss of MHCII in stromal cells or fibroblasts elevated metastatic tumor growth, whereas co-implantation of cancer cells and MHCII-positive fibroblasts, in contrast to MHCII-deficient fibroblasts, reduced tumor progression. Together, these findings establish immune-stromal crosstalk that links CD4⁺ T cell activation to a fibroblast-differentiation program associated with enhanced control of pancreatic cancer.

The responsiveness of fibroblast MHCII expression to immune signals highlights the plasticity of the CAF compartment. We and others have observed changes in CAF composition within the tumor microenvironment following combined PD-1 blockade and CCR8-or CTLA-4-directed intratumoral Treg-depleting therapies^25,28^. In our study, MHCII expression increased in both Ly6C-positive and Ly6C-negative CAFs, suggesting that the MHCII^+^CAF phenotype is an inducible state accessible to distinct fibroblast populations. This phenotypic flexibility, together with evidence of dual cellular origins of antigen-presenting CAFs^14,24^, supports a model in which both cellular origin and local immune signals shape the MHCII⁺CAF compartment.

An important question is whether MHCII expression merely marks an immune-responsive fibroblast state or contributes functionally to tumor restraint. Previous studies have linked MHCII⁺CAF abundance to patient survival and therapeutic responses^14,28,48,49^, but these associations do not establish a functional requirement for MHCII expression in CAFs. We therefore combined bone marrow chimera experiments, fibroblast-specific MHCII deletion, and fibroblast–cancer cell co-implantation studies to assess its contribution to tumor control. MHCII deficiency in the non-hematopoietic compartment increased metastatic burden. Fibroblast-specific MHCII deletion also promoted metastases following Treg ablation, consistent with MHCII expression in fibroblasts promoting enhanced antitumor immunity^50–52^. In addition, co-implantation of cancer cells with MHCII-sufficient fibroblasts restrained primary and metastatic tumor progression compared with co-implantation with MHCII-deficient fibroblasts. Together, these complementary approaches establish that MHCII expression in CAFs is both necessary and sufficient for tumor control in our PDAC models

The antitumor activity of MHCII^+^CAFs may involve positive feedback with CD4^+^ T cells. Although antigen presentation by non-hematopoietic cells is often associated with tolerance in healthy tissues, these cells can also activate CD4^+^ T cells during inflammation. For example, lung fibroblasts can support CD4⁺ T cell priming during bacterial infection^53^, and MHCII⁺ intestinal epithelial cells promote CD4⁺ T cell responses in graft-versus-host disease^54^. Within tumors, Th1-derived IFNγ induces MHCII expression in fibroblasts, and MHCII^+^CAFs could, in turn, sustain local CD4^+^ T cell responses that help control tumor progression^25,27^. Beyond cognate antigen recognition, interactions with MHC class II molecules can support CD4^+^ T cell metabolic fitness and survival^55^, further illustrating how MHCII^+^CAFs could sustain an immune-supportive niche.

Changes in the balance between CAF subsets may also contribute to tumor control. In our models, increased MHCII⁺CAF frequency coincided with a reduction in myCAFs, consistent with previous evidence of stromal remodeling following Treg ablation^39^. Previous studies have shown that myCAFs can promote tumor progression by supporting cancer cell proliferation and invasion^2,3^, while interventions that promote fibroblast quiescence can limit tumor progression^50–52^. Together, these observations suggest that an increase in MHCII^+^CAFs could help restrain PDAC progression by reinforcing CD4⁺ T cell responses and shifting the balance of CAF functions toward tumor restraint.

Our findings also broaden the mechanisms through which conventional CD4⁺ T cells coordinate antitumor responses in the absence of Tregs. CD4⁺ T cells have been shown to promote antitumor responses through interactions with dendritic cells^34,56^, NK cells^39^, and myeloid cells^57^. Our study identifies CAFs as an additional component of this response, extending the effects of conventional CD4⁺ T cell activation to the tumor stroma through IFNγ signaling. IFNγ promotes antitumor immunity by enhancing antigen presentation, inducing M1 macrophage polarization, and supporting cytotoxic T cell responses, yet its effects vary with cellular context^58^. In pancreatic cancer cells, prolonged IFNγ exposure has been reported to promote TAP2 ubiquitination, leading to reduced MHC class I expression and impaired CD8^+^ T cell recognition^59^. IFNγ-induced MHCII^+^CAF reprogramming may therefore provide a complementary mechanism for tumor control when CD8^+^ T cell responses are compromised. Supporting this hypothesis, tumor control following Treg ablation involved conventional CD4^+^ T cells and MHCII^+^CAFs but did not require CD8^+^ T cells. IFNγ neutralization, however, abrogated both tumor control and MHCII^+^CAF reprogramming. Collectively, these findings support a CD4^+^ T cell-IFNγ-CAF axis that enables tumor restraint without requiring CD8^+^ T cells and highlight MHCII^+^CAFs as an additional compartment through which immune activation can regulate tumor progression.

Our mechanistic findings suggest a promising therapeutic opportunity, supported by our observation that combined CCR8-directed Treg targeting and PD-1 blockade increased MHCII⁺CAF abundance alongside tumor control. These results demonstrate that the immune–stromal response identified through Treg ablation can also be engaged through a therapeutically relevant intervention. Targeting Treg-mediated suppression may therefore offer a practical strategy to enhance conventional CD4⁺ T cell responses while promoting tumor-restraining CAF functions. Stromal therapies that reduce immune exclusion or suppressive signaling while preserving MHCII⁺CAFs could complement this approach by supporting CD4⁺ T cell access and activity. Our study provides a mechanistic basis for IFNγ-dependent MHCII⁺CAFs supporting antitumor immunity and justifies the development of strategies to increase MHCII expression on CAFs for the immunotherapeutic treatment of PDAC.

## EXPERIMENTAL MODEL AND SUBJECT DETAILS

### Cell lines

All murine pancreatic cancer cell lines were kindly provided by Dr. Monte Winslow (Stanford University). 6421c2, 6694c2, and 6419c5, derived from KPCY (Kras^LSL-G12D/+^;Trp53^LSL-R172H/+^;Pdx1-Cre;Rosa26^YFP/YFP^) mice on a c57BL/6 background, originated from Dr. Ben Stanger’s Laboratory (University of Pennsylvania)^29^. FC1245, derived from KPC (Kras^LSL-G12D/+^;Trp53^LSL-R172H/+^;Pdx1-Cre) mice on a c57BL/6 background, originated from Dr. David Tuveson’s Laboratory (Cold Spring Harbor Laboratory)^30^. All cancer cell lines were maintained in Dulbecco’s Modified Eagle Medium (DMEM, Gibco) supplemented with 10% fetal bovine serum (Biowest), 1% penicillin-streptomycin (Gibco), and 1% L-Glutamine (Gibco). Primary fibroblasts were isolated from the lungs of wild-type c57BL/6 or *MHCII^-/-^*mice and maintained in DMEM supplemented with 20% FBS, 1% P/S, 1% L-Glu, and 0.1% Plasmocin Prophylactic (Invitrogen). All cells were grown at 37°C with 5% CO_2_. The cell lines were tested for Mycoplasma by PCR and were maintained in culture for a maximum of six passages between thawing and use in experiments.

### In vivo animal studies

All mice used were bred onto a C57BL/6 background. Littermates or age-matched control mice (7-20 weeks old) were used in all the mouse experiments. The *Foxp3^GFP-DTR^* mice express human diphtheria toxin receptor (DTR) and EGFP genes from the Foxp3 locus without disrupting expression of the endogenous Foxp3 gene^60^. These mice were a gift from Dr. Alexander Rudensky (Memorial Sloan Kettering Institute; JAX #016958). The *MHCII^-/-^* (*I-A^b-/-^*) mice (JAX #003584), *Rag2^-/-^* mice (JAX #008449), and CD45.1^+^c57BL/6 mice (JAX #033076) were obtained from The Jackson Laboratory. The *Pdgfrα^CreERT^*^2^*^/+^;I- Ab^fl/fl^* mice were generated by crossing the *Pdgfrα^CreERT^*^2^*^/+^* mice (JAX #032770) with the *I-Ab^fl/fl^* mice (JAX #013181) in our laboratory, both of which were obtained from The Jackson Laboratory. All the experiments were conducted in accordance with the Institutional Animal Care and Use Committee guidelines of the University of California, Berkeley (AUP-2017-05-9915-3).

## METHOD DETAILS

### Orthotopic models

For orthotopic models of PDAC, 6421c2, 6694c5, 6419c2, or FC1245 (2 x 10^5^ cells) prepared in 50μL DPBS (Gibco) were injected at the tail end of the pancreas in 7-to-12-week-old C57BL/6 background mice. For immune and stromal phenotyping studies, tumors were harvested on day 16 post-inoculation. For the co-implantation model, 1x10^5^ 6421c2 and 5x10^5^ primary fibroblasts in 50μL DPBS were injected at the tail end of the pancreas. The co-implanted tumors were harvested on day 17 post-inoculation. For diphtheria toxin (DT, EMD Millipore) treatments, mice received vehicle control (HBSS, Gibco) or 1 μg DT per mouse intraperitoneally (i.p.) every other day starting on day 7 post-tumor inoculation. For tamoxifen (TAM, Sigma) treatment, *Pdgfrα^CreERT^*^2^*;I-Ab^fl/fl^* mice and the CreERT2-negative littermates received 4mg tamoxifen via oral gavage (p.o.) every 4 days starting on day 5 before tumor inoculation. Tumors were harvested on day 21 post-inoculation. Both DT and tamoxifen treatments were given continuously at the frequency stated above throughout the tumor model.

### In vivo antibody-mediated cell depletion

For T cell depletion, rat IgG2b (Clone LTF-2, Leinco Technologies), anti-mouse CD4 antibody (Clone GK1.5, Leinco Technologies) or anti-mouse CD8 antibody (Clone 53-5.8, Leinco Technologies) at 200 μg/mouse were administered i.p. on day 1 pre-tumor implantation and days 1, 7, and 13 post-implantation. For IFNγ neutralization, rat IgG1 (Clone GL113, Leinco Technologies) or anti-mouse IFNγ (Clone XMG1.2, BioXcell) was administered at 200μg/mouse i.p. on days 5, 8, 11, and 14 post-inoculation. For αCCR8 and αPD-1 treatment, mouse IgG2a (Clone C1.18.4, BioXcell), anti-mouse CCR8 (Gilead Sciences), anti-mouse PD-1 (Gilead Sciences), or a combination of both antibodies was administered i.p. on days 5, 8, 11, and 14 post-implantation (200μg per mouse per injection). All tumors were harvested on day 16 post-tumor inoculation.

### Bone marrow chimera generation

For the stromal MHCII-deficient study, the host wild-type C57BL/6 and MHCII^KO^ (*I-A^b-/-^*) mice were given two doses of 5 Gy irradiation, 16 hours apart. An XRad320 (Precision X-Ray Irradiation) was used to administer the irradiation. Four hours after the second irradiation treatment, 1 × 10^7^ bone marrow cells from the wild-type C57/BL6 mice were transferred to the host mice via tail-vein injection to reconstitute MHCII expression in hematopoietic cells. The bone marrows from femurs, tibias, and humeri were harvested from the donor mice and flushed with Opti-MEM (Gibco) using a 27-gauge syringe, followed by red blood cell removal using Ammonium-Chloride-Potassium (ACK) lysis buffer (150 mmol/L of NH_4_Cl, 10 mmol/L of KHCO_3_, 0.1 mmol/L of Na_2_EDTA, pH 7.3). The bone marrow was washed with PBS twice prior to tail-vein injection. The host mice receiving bone marrow transfer were administered water supplemented with sulfamethoxazole-trimethoprim oral suspension (ANI Pharmaceuticals) for 30 days and were monitored daily. The bone marrow was permitted to reconstitute recipient mice for 60 days. Gender-matched CD4^+^ T cells from CD45.1^+^ C57BL/6 donor mice were transferred to the reconstituted recipient mice at a one-mouse-to-mouse transfer (approximate 1.5 × 10^7^ cells) to rebuild the CD4 repertoire in MHCII-deficient hosts. Four days post-CD4 transfer, 2 × 10^5^ 6421c2 cells were implanted orthotopically. Tumors were harvested on day 21 post-implantation.

For fibroblast conditional knockout of MHCII, bone marrow chimeras were generated as described above. Specifically, 1 × 10^7^ bone marrows from *Foxp3^DTR^* mice were transferred to the irradiated recipient mice, *Pdgfrα^CreERT^*^2^*^/+^;I-Ab^fl/fl^* and *Pdgfrα^+/+^;I-Ab^fl/fl^*. Following reconstitution, 2 × 10^5^ 6421c2 cells were implanted orthotopically. Tamoxifen (4 mg, p.o.) was given every 4 days starting on day 5 before tumor inoculation. DT (1 μg, i.p.) was given every other day starting on day 7 post-implantation. Tumors were harvested on day 21 post-implantation.

### Primary fibroblast isolation and preparation

Primary fibroblasts were isolated from the lungs of wild-type c57BL/6 or MHCII^-/-^(*I-A^b-/-^*) mice using adjusted protocols as follows^46^. Lungs were chopped into small pieces (< 2 mm in length) and seeded onto T75 flasks in DMEM supplemented with 20% FBS, 1% P/S, 1% L-Glu, and 0.1% Plasmocin Prophylactic (Invitrogen). On day 3, the primary culture was replenished with fresh culture media. On day 10 post-harvest, cells were trypsinized (Gibco) and filtered through 40μm cell strainer (Falcon) to isolate primary fibroblasts. Fibroblasts were maintained in culture up to day 14 post-harvest. Before tumor inoculation in the co-implantation study, primary fibroblasts were treated with 20ng/mL IFNγ for 24h, followed by two PBS washes.

### Murine lymphocyte and stromal cell isolation

Pancreatic tumors were resected and minced into 1 mm^3^ isolates, then digested in RPMI media supplemented with HEPES, 20 μg/mL of DNase I (Roche), 0.8 mg/mL of Collagenase P (Roche), and 0.02% bovine serum albumin (BSA, VWR) on an orbital shaker for 30 minutes at 37°C. All the cell suspensions were passed through 70 μm-and then 40 μm-filters to generate single-cell suspensions. Tumor cell suspensions were subjected to red blood cell lysis using ACK lysis buffer (150 mmol/L of NH_4_Cl, 10 mmol/L of KHCO_3_, 0.1 mmol/L of Na_2_EDTA, pH 7.3) and resuspended in ice-cold FACS buffer (PBS supplemented with 2 mmol/L of EDTA and 2% FBS) before cell staining or in vitro stimulation. Single-cell suspensions of cervical and mesenteric lymph nodes were generated through mechanical disruption of harvested lymph nodes and were passed through 70μm- and then 35μm-filters. Cell suspensions were resuspended in ice-cold FACS buffer before cell staining or in vitro stimulation. Cytokine staining was performed on 1 × 10^6^ cells after 4 hours of in vitro stimulation in Opti-MEM media supplemented with Brefeldin A (eBioscience), 10 ng/mL of phorbol 12-myristate 13-acetate (Sigma), and 0.25 μmol/L of ionomycin (Sigma) at 37°C. Pro-inflammatory cytokine ELISA was performed on supernatant of 1 × 10^6^ cells after 4 hours of in vitro culture in Opti-MEM media at 37°C.

### Flow cytometry

Flow cytometry was performed on a BD LSR Fortessa X20 (BD Biosciences) or a BD FACSympohny A3 Cell Analyzer (BD Biosciences). Datasets were analyzed using FlowJo software (Tree Star). All samples analyzed were reconstituted in single-cell suspensions using ice-cold PBS, and Fc blocked using TruStain FcX Plus Antibody (BioLegend; RRID: AB_2783138) for 30 minutes at 4°C prior to staining. Dead cells were stained with Live/Dead Fixable Blue or Aqua Dead Cell Stain Kit (Molecular Probes) in PBS for 30 minutes at 4°C. Cell surface proteins were stained for 30 minutes at 4°C using a mixture of fluorophore-conjugated antibodies. Surface marker stains for murine samples were carried out with anti-mouse CD3 (17A2, BD Biosciences; RRID: AB_2738781; BioLegend; RRID: AB_1595492), anti-mouse CD4 (RM4-5, BD Biosciences; RRID: AB_2870165), anti-mouse CD8α (53-6.7, BD Biosciences; RRID: AB_2870090; BioLegend; RRID: AB_2562610), anti-mouse CD11b (M1/80, BioLegend; RRID: AB_2561373), anti-mouse CD11c (N418, BioLegend; RRID: AB_313779), anti-mouse CD25 (PC61, BioLegend; RRID: AB_312857 & AB_893288), anti-mouse CD31 (390, BioLegend; RRID: AB_830756), CD44 (IM7, Thermo; RRID: AB_1272244), CD45 (30-F11, BD Biosciences; RRID: AB_2651134), anti-mouse CD45.1 (A20, BioLegend; RRID: AB_2563379), anti-mouse CD45.2 (104, BioLegend; RRID: AB_313445), CD45R (RA3, BD Biosciences; RRID: AB_2738470), anti-mouse CD62L (MEL-14, BioLegend; RRID: AB_313099 & AB_493380), anti-mouse CD86 (GL-1, BD Biosciences; RRID: AB_2244452), anti-mouse CD90.2 (30-H12, BD Biosciences; RRID: AB_10613293), anti-mouse CD103 (2E7, BD Biosciences; RRID: AB_1227502 & AB_2629724), anti-mouse CCR8 (SA214G2, BioLegend; RRID: AB_2728199), anti-mouse EpCAM (G8.8, BioLegend; RRID: AB_1134099), anti-mouse F4/80 (BM8, BioLegend; RRID: AB_893478), anti-mouse Ly6C (HK1.4, BioLegend; RRID: AB_2562351 & AB_1186133), anti-mouse Ly6G (1A8, BioLegend; RRID: AB_2562567 & AB_2566317), anti-mouse MHC Class I H2-K^b^/H2-D^b^ (28-8-6, BioLegend; RRID: AB_2750194), anti-mouse MHC Class II I-A/I-E (M5/114.15.2, BioLegend; RRID: AB_2565975), anti-mouse NKG2A/C/E (20d5, BD Biosciences; RRID: AB_2738782), anti-mouse NKp46 (29A1.4, BioLegend; RRID: AB_10552741), anti-mouse PD-1 (29F.1A12, BioLegend; RRID: AB_2566158. RMP1-30, BioLegend; RRID: AB_572017), anti-mouse PDPN (8.1.1, BioLegend; RRID: AB_2814017). For experiments where no intracellular staining was required, cells were fixed in 2% paraformaldehyde for 10 minutes at 4°C. For experiments requiring intracellular staining, cells were fixed using the Tonbo Foxp3 Transcription Factor Staining Buffer Kit (Tonbo Bioscience) for 30 minutes at 4°C prior to intracellular staining. Intracellular staining was performed using anti-mouse Foxp3 (FJK-16S, Thermo; RRID: AB_465243), anti-mouse GATA3 (16E10A23, BioLegend; RRID: AB_2563217), anti-mouse IFNγ (XMG1.2, BioLegend; RRID: AB_893526), anti-mouse Ki67 (B56, BD Biosciences; RRID: AB_10611874), anti-mouse RORγt (Q31-378, BD Biosciences; RRID: AB_2651150), anti-mouse T-bet (4B10, BioLegend; RRID: AB_2561761) for overnight at 4°C. Post-staining, cells were resuspended in PBS and filtered through a 35 μm nylon mesh (Fisher Scientific) before data acquisition.

### Bulk RNA-sequencing

Orthotopic 642c2 tumors treated with HBSS (vehicle) or DT were processed into single-cell suspensions and sorted on the BD FACSAira Fusion Cell Sorter (BD Biosciences). Prior to sorting, cell suspensions were Fc blocked using TruStain FcX Plus Antibody for 30 minutes at 4°C, followed by viability staining with Propidium iodine (Sigma) in PBS for 30 minutes at 4°C. Cell surface proteins were stained for 30 minutes at 4°C using a mixture of fluorophore-conjugated anti-mouse antibodies: anti-CD4 (GK1.5, BioLegend; RRID AB_312707), anti-CD8α (53-6.7, BioLegend; RRID AB_2562609), anti-CD25 (PC61, BioLegend; RRID AB_893288), anti-CD31 (390, BioLegend; RRID AB_493498), anti-CD45 (30-F11, BioLegend; RRID AB_2561392), anti-mouse EpCAM (G8.8, BioLegend; RRID: AB_1134099), anti-mouse Ly6C (HK1.4, BioLegend; RRID: AB_1186133), anti-mouse MHC Class II I-A/I-E (M5/114.15.2, BioLegend; RRID: AB_1659252), and anti-mouse PDPN (8.1.1, BioLegend; RRID: AB_2814017). Four CAF subsets, Ly6C^-^MHCII^-^(myCAF), Ly6C^+^MHCII^-^(iCAF), Ly6C^-^MHCII^+^CAF, and Ly6C^+^MHCII^+^CAF, and two CD4 subsets, CD25^-^Foxp-DTR-GFP^-^(Tconv) and CD25^+^Foxp-DTR-GFP^+^ (Treg), were sorted for bulk RNA-sequencing. The sorted cells were pelleted at 800 xg for 8 min minutes at 4°C, then subjected to RNA isolation with RNeasy Micro Kit (Qiagen) following the manufacturer’s protocol. Bulk RNA-sequencing library preparations were performed by Signios Bio using Takara SMART-Seq mRNA Library Prep (Takara Bio) followed by sequencing on the NovaSeq paired-end 150 bp (Illumina).

### Immunohistochemistry and immunofluorescence

Pancreatic tumors were resected, embedded in Tissue-Plus O.C.T. Compound (Fisher), and flash-frozen in liquid nitrogen vapor. Frozen tissue blocks were sectioned at 8-15 μm using a Thermo NX50 Cryostat Microtome (Thermo Scientific). Trichrome staining was performed on frozen sections using the VitroView Masson’s Trichrome Stain Kit following the manufacturer’s protocol. Histology images were taken using Zeiss Axioscan 7 microscope (Zeiss). For immunofluorescence (IF) staining, frozen tumor sections were fixed in 4% paraformaldehyde for 15 minutes at room temperature, washed with PBS 3 times, and blocked with 1% BSA in PBS for 1 hour at room temperature. Tissue sections were then stained with rabbit anti-mouse αSMA (Cell Signaling, Catalog 19245T, 1:200) for 2 hours, followed by incubation with Donkey anti-Rabbit IgG (H+L) Highly Cross-Adsorbed Secondary Antibody, Alexa Fluor 546 (Thermo Fisher, Catalog A10040, 1:500) for 1 hour in the dark at room temperature. Stained tissue sections were mounted with ProLong Gold Antifade Mountant with DNA stain DAPI (Fisher Scientific, Catalog P36931). IF Images were taken using a Zeiss LSM 880 NLO Airyscan microscope (Zeiss). Imaging analysis was performed using QuPath and ImageJ software.

### ELISA

Tumors from mice receiving HBSS (vehicle), DT + IgG2b, or DT + anti-CD4 antibody injections were dissected on day 16 post-tumor implantation. One million cells were plated in 200 μL of Opti-MEM (Gibco) in a 96-well plate and incubated at 37°C for 4 hours. Supernatants were collected, and ELISA was performed using LEGENDplex Mouse Inflammation Panel, 13-plex (BioLegend, Catalog 740446) following the manufacturer’s protocol.

### Quantification and statistical analysis

The p-values were obtained from unpaired two-tailed Student’s t-tests for all statistical comparisons between two groups, and data were displayed as mean ± SEM. For multiple comparisons, one-way ANOVA with Tukey’s multiple comparisons test was used. For multi-group comparisons, two-way ANOVA with Šidák’s multiple comparison test was used. Survival analysis was performed using the Mantel-Cox Logrank test. The p-values are denoted in figures by *p<0.05, **p<0.01, ***p<0.001, and ****p<0.001

## Acknowledgements

We thank Kartoosh Heydari, Harman Dhaliwal, and Melaine Delcroix of the UC Berkeley Cancer Research Laboratory Flow Cytometry Facility. We thank Holly Aaron, Benjamin Smith, Cherise Stanley, and Luis Alvarez of the UC Berkeley Cancer Research Laboratory Molecular Imaging Center. We thank Signios Bioscience for their technical support and RNA sequencing service. The authors would also like to thank all members of the DuPage laboratory for providing guidance on the research approach. This work was supported by the National Institutes of Health DP2CA247830 and R01CA292818 (to M. DuPage), R01DE027750 (to J.E. Elias & J.B. Sunwoo), K99CA304503 (to J. Hung), AI100829 (T32 to J. Hung), the California Institute of Generative Medicine Grant EDUC4-12790 (to J. Hung), Gilead Sciences (to M. DuPage), a Cancer Research Coordinating Committee fellowship (to S. Silveria-Davies), and the UC President’s Pre-Professoriate Fellowship (to B.G. Maker). M. DuPage was also supported as a Pew-Stewart Scholar and as a St. Baldrick’s Scholar with generous support from Hope with Hazel.

---

**Supplementary Figure 1.**
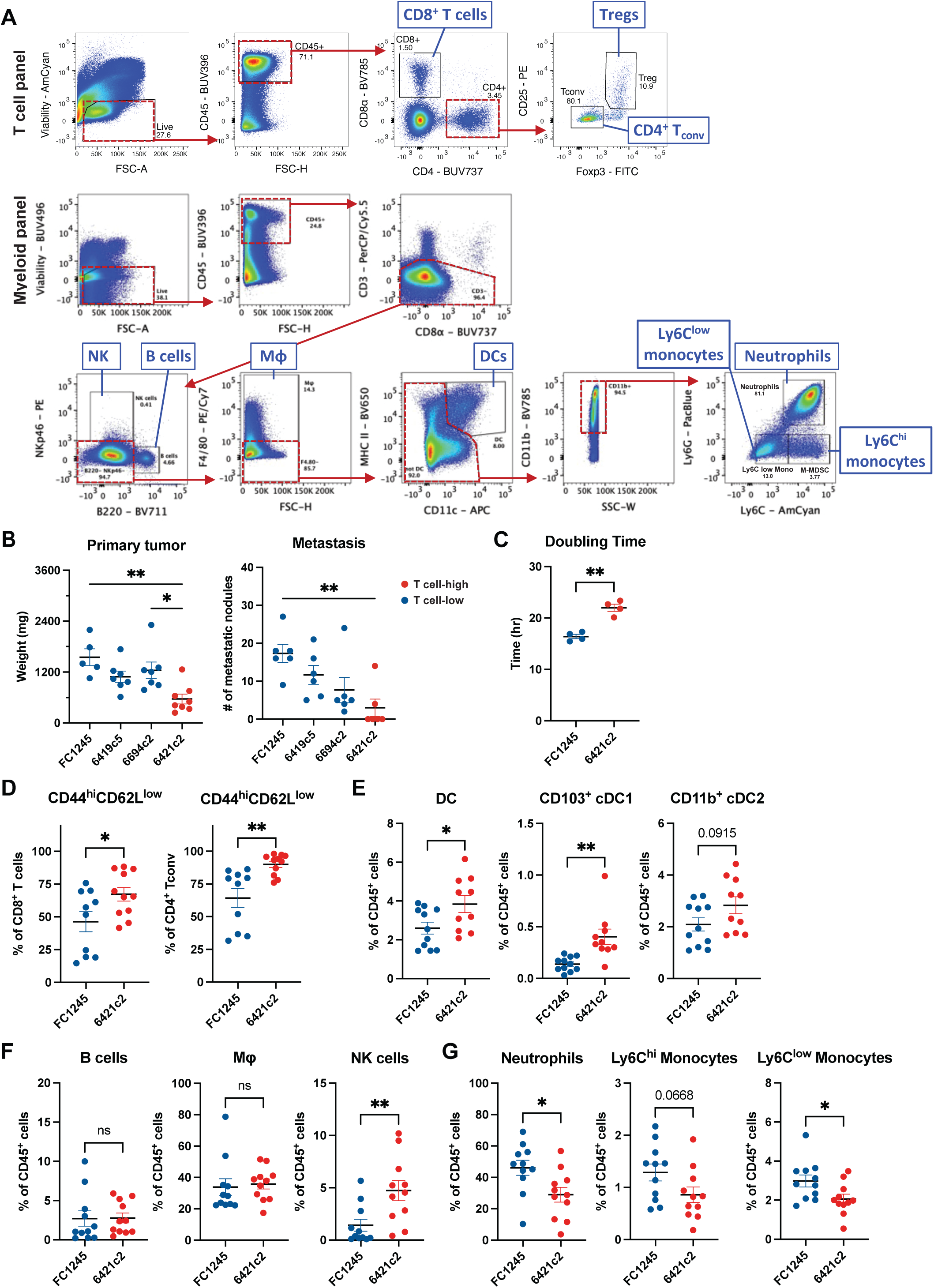
Distinct immune infiltrates in orthotopic tumors derived from four pancreatic cancer cell lines. (A) Gating strategies for T cell and myeloid cell panels. (B) Primary tumor mass and number of metastatic nodules on day 19 post-tumor implantation. (C) In vitro doubling times of FC1245 and 6421c2 cells. (D) Frequencies of CD44^hi^CD62^low^ cells among CD8^+^ and CD4^+^ T cells in FC1245 and 6421c2 tumors on day 16 post-implantation. Frequencies of (E) DCs, cDC1, cDC2, (F) B cells, macrophages, NK cells, (G) neutrophils, and monocytes among CD45^+^ cells in FC1245 and 6421c2 tumors. (B, D-G), data are pooled from two independent experiments. Data are shown as mean ± SEM; *p<0.05; **p<0.01. (B), One-way ANOVA with Tukey’s multiple comparison test. (C-F), unpaired Student’s t-test.

**Supplementary Figure 2.**
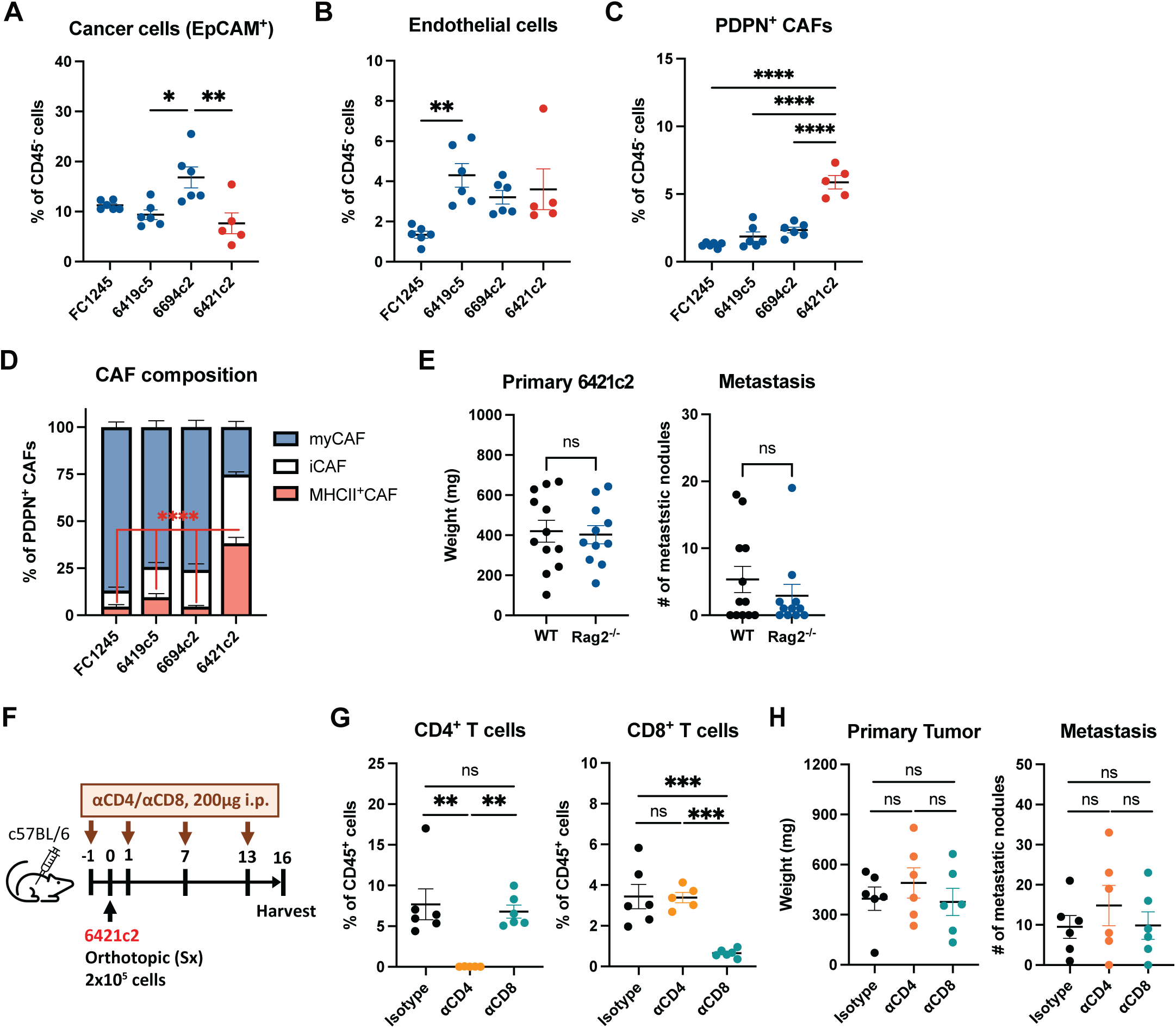
Distinct stromal compartments in pancreatic tumors with different T cell infiltrates. Frequencies of (A) cancer cells, (B) endothelial cells, and (C) PDPN^+^CAFs among CD45^-^cells in orthotopic tumors derived from four PDAC cell lines on day 16 post-implantation. (D) Frequencies of CAF subsets among PDPN^+^CAFs in four PDAC tumors. (E) Primary tumor mass and number of metastatic nodules in 6421c2-bearing wild-type and Rag^-/-^mice on day 16 post-implantation. (F) Schematic of IgG2b, CD4, or CD8 depletion in 6421c2-bearing mice. Tumors were harvested on day 16 post-implantation. (G) Frequencies of CD4^+^ and CD8^+^ T cells among CD45^+^ cells in 6421c2 tumors at harvest. (H) Primary tumor mass and number of metastatic nodules in 6421c2 tumors treated with IgG2b, anti-CD4, or anti-CD8 antibodies. (A-E), Data are pooled from two independent experiments. (G, H), Data shown are representative of two independent experiments with the same results. Data are shown as mean ± SEM; *p<0.05; **p<0.01; ***p<0.001; ****p<0.0001. (A-C, G, H), One-way ANOVA with Tukey’s multiple comparison test. (D), Ordinary Two-way ANOVA with Šídák’s multiple comparison test. (E, F), unpaired Student’s t-test.

**Supplementary Figure 3.**
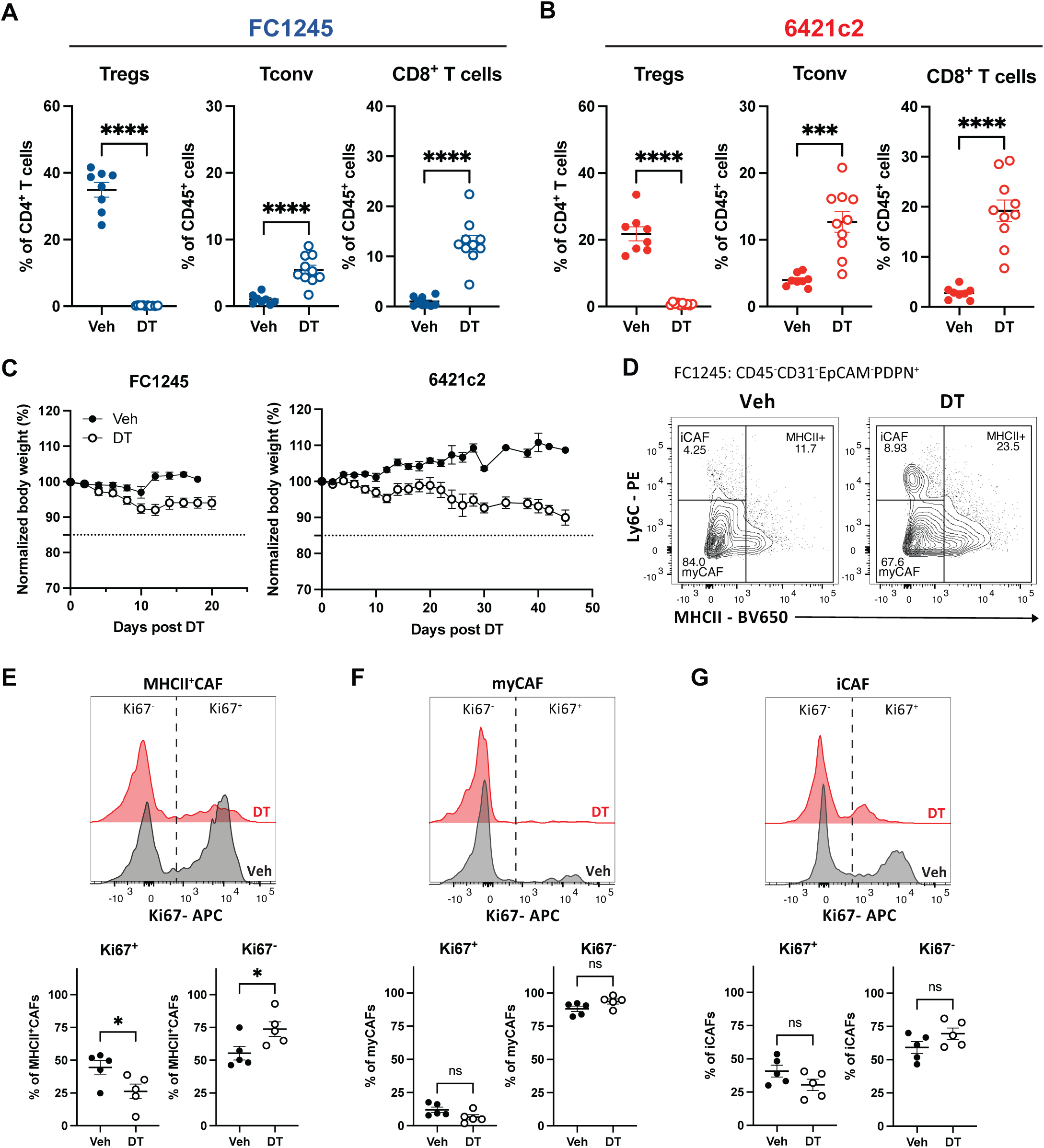
Treg ablation changes T cell and CAF subsets in FC1245 and 6421c2 tumors. Frequencies of Tregs among CD4^+^ T cells and frequencies of conventional CD4^+^ T cells and CD8^+^ T cells among CD45^+^ cells in (A) FC1245 and (B) 6421c2 tumors treated with vehicle or DT injections. (C) Body weight of FC1245-and 6421c2-bearing mice treated with repeated vehicle or DT injections, every other day, starting on day 7 post-tumor implantation. (D) Representative flow plots of CAF subsets in FC1245 tumors treated with vehicle or DT injections. Representative histograms and quantification of Ki67 staining in (E) MHCII^+^CAFs, (F) myCAFs, and (G) iCAFs in 6421c2 tumors treated with vehicle or DT injections. Pre-gates for the CAF histogram: CAF subsets defined by Ly6C and MHCII from CD45^-^CD31^-^EpCAM^-^PDPN^+^ cells. (A-C), Data are pooled from two independent experiments. (E-G), Data shown are representative of two independent experiments with the same results. Data are shown as mean ± SEM; *p<0.05; ***p<0.001; ****p<0.0001. (A, B, E), unpaired Student’s t-test.

**Supplementary Figure 4.**
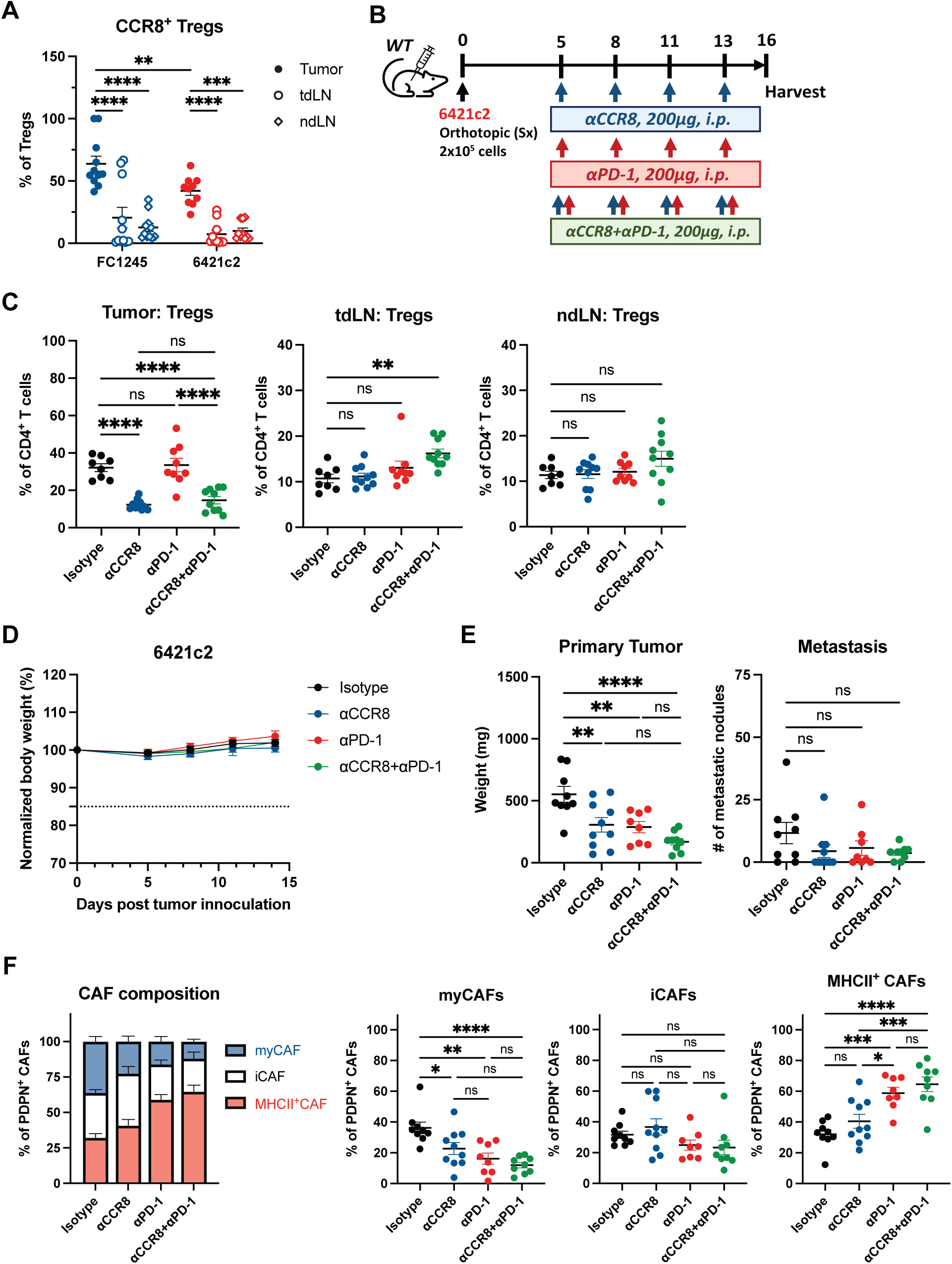
Combination therapy targeting CCR8 and PD-1 promotes tumor control and upregulation of MHCII^+^CAFs. (A) Frequencies of CCR8^+^Tregs among all Tregs in tumors, tumor-draining lymph nodes, and non-draining lymph nodes of FC1245 and 6421c2 orthotopic models. (B) Schematic of IgG2a, anti-CCR8, anti-PD-1, or anti-CCR8+anti-PD-1 treatment in 6421c2-bearing mice. Tumors were harvested on day 16 post-implantation. (C) Frequencies of Tregs among CD4^+^ T cells in tumors, tumor-draining lymph nodes, and non-draining lymph nodes of 6421c2-bearing mice receiving IgG2a, anti-CCR8, anti-PD-1, or combined antibody treatment. (D) Mouse body weight during the immunotherapy treatment. (E) Primary tumor mass and number of metastatic nodules in the 6421c2 model receiving anti-CCR8, anti-PD-1, or combined antibody treatment. (F) Frequencies of CAF subsets and number per gram of tumor in PDPN^+^CAFs in the 6421c2 model receiving anti-CCR8, anti-PD-1, or combined treatment. Data are pooled from two independent experiments. Data are shown as mean ± SEM; *p<0.05; **p<0.01; ***p<0.001; ****p<0.0001. (C, E), One-way ANOVA with Tukey’s multiple comparison test. (A, F), Ordinary Two-way ANOVA with Šídák’s multiple comparison test.

**Supplementary Figure 5.**
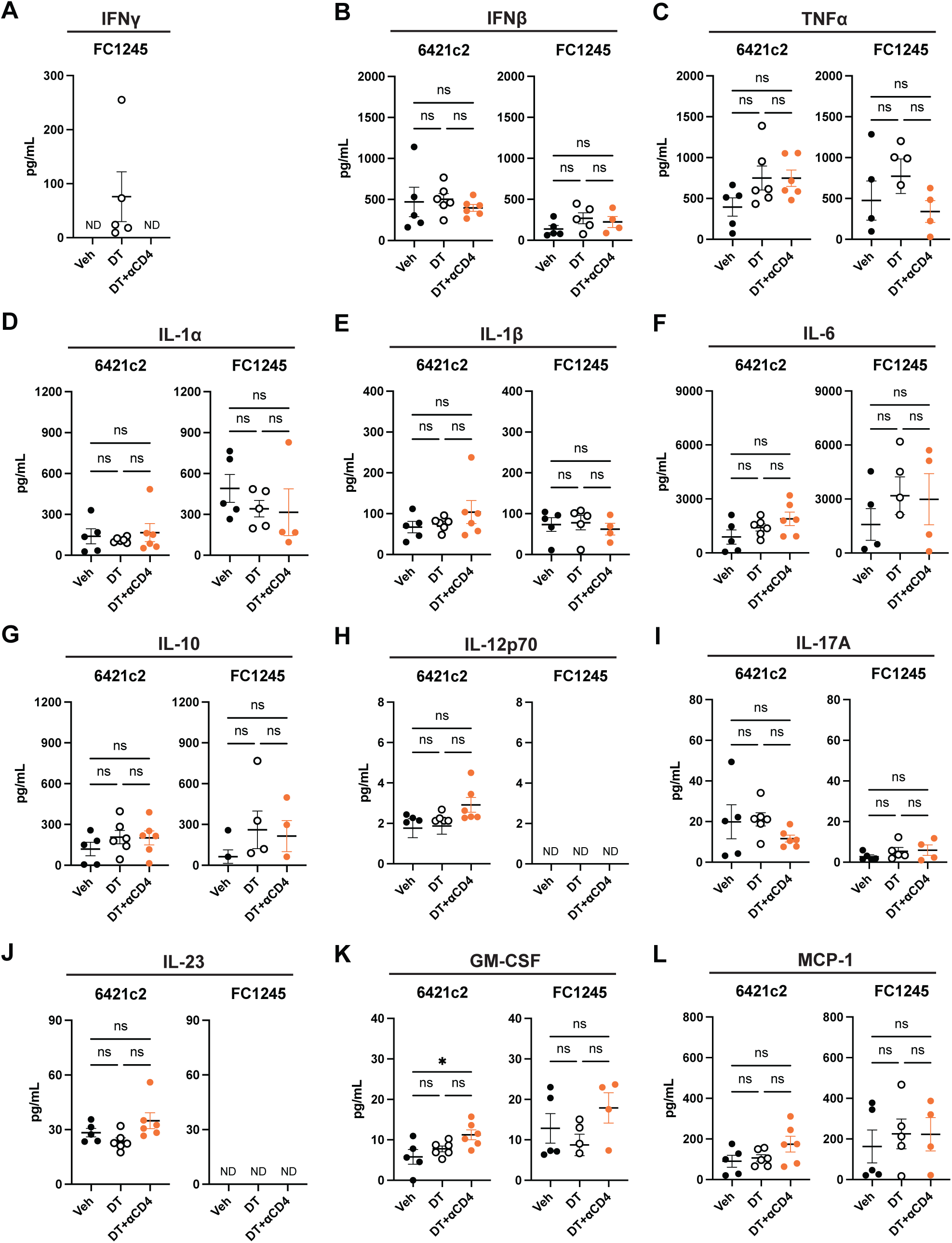
Characterization of proinflammatory cytokines in Treg-ablated tumors. (A) IFNγ concentration in FC1245 tumors treated with vehicle, DT + IgG2b, or DT + anti-CD4 antibodies. Concentrations of (B) IFNβ, (C) TNFα, (D) IL-1α, (E) IL-1β, (F) IL-6, (G) IL-10, (H) IL-12p70, (I) IL-17A, (J) IL-23, (K) GM-CSF, and (L) MCP-1 in 6421c2 and FC1245 tumors treated with vehicle, DT + IgG2b, and DT + anti-CD4 antibodies. Data are shown as mean ± SEM; ND, not detectable. Statistics were performed using One-way ANOVA with Tukey’s multiple comparison test. IL-27 was also measured but remained undetectable regardless of Treg ablation or CD4 depletion.

**Supplementary Figure 6.**
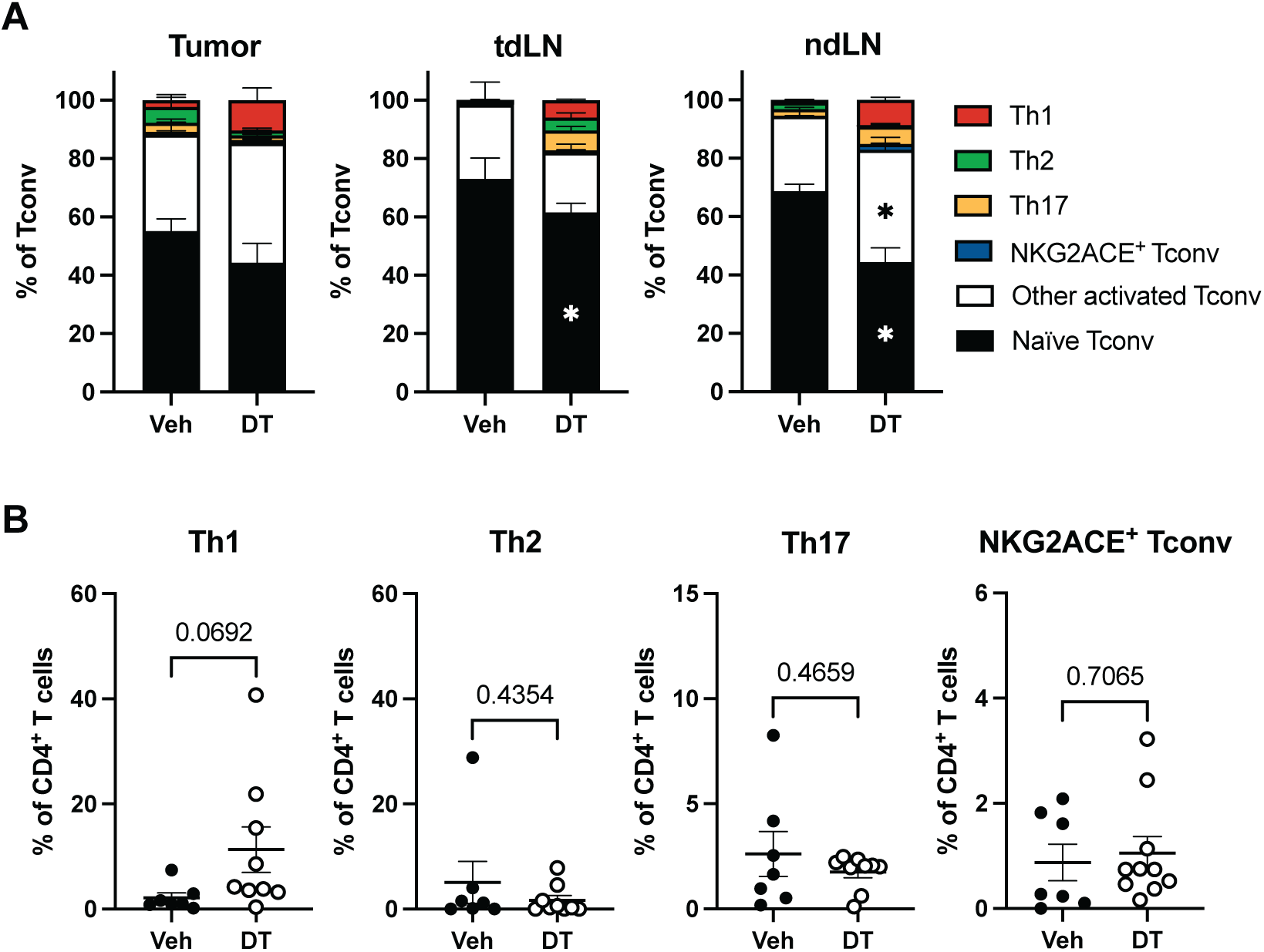
Characterization of conventional CD4^+^ T cell subsets in Treg-ablated FC1245 tumors. (A) Composition of conventional CD4^+^ T cell subsets in tumor, tumor-draining lymph nodes, and non-draining lymph nodes in the FC1245-bearing mice treated with vehicle or DT. (B) Frequencies of Th1(T-bet^+^), Th2(GATA3^+^), Th17(RORγt^+^), and NKG2ACE^+^ Tconv among CD4^+^ T cells in FC1245 +/-DT. Data are pooled from two independent experiments. Data are shown as mean ± SEM; *p<0.05. (A), Ordinary Two-way ANOVA with Šídák’s multiple comparison test. (B), unpaired Student’s t-test.

**Supplementary Figure 7.**
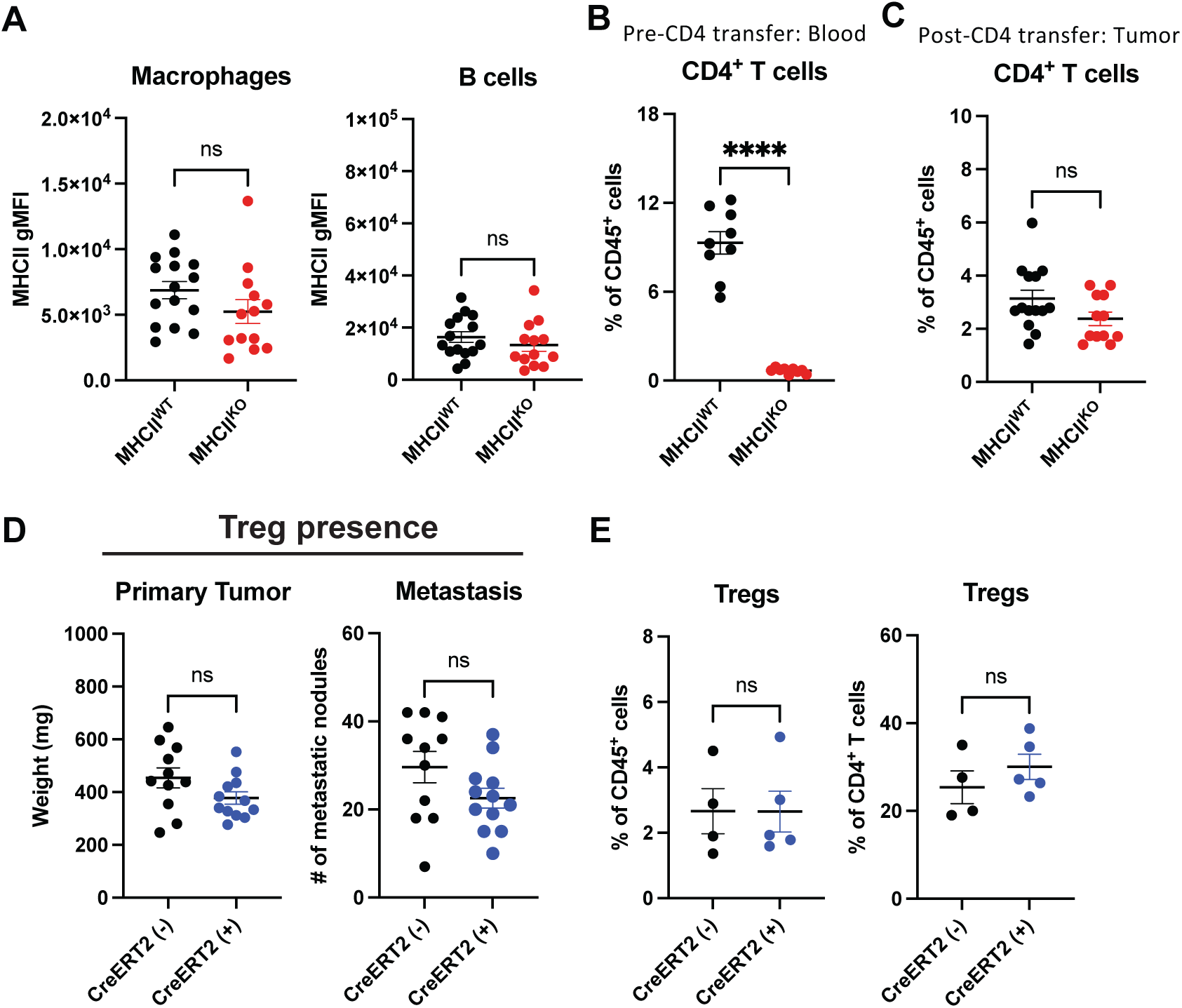
Validation of stromal and fibroblast-specific MHCII-deficiency models. (A) MHCII gMFI in macrophages and B cells in tumors of the stromal MHCII deficiency model at harvest. (B) Frequencies of CD4^+^ T cells among CD45^+^ cells in the blood of *MHCII^WT^* and *MHCII^KO^* recipient mice before CD4^+^ T cell adoptive transfer. (C) Frequencies of CD4^+^ T cells among CD45^+^ cells in 6421c2 tumors of *MHCII^WT^*and *MHCII^KO^* recipient mice at harvest. (D) Primary tumor mass and number of metastatic nodules in 6421c2 tumors with MHCII^WT^ (CreERT2-negative) or MHCII^KO^ (CreERT2-positive) CAFs following HBSS (vehicle) injections. (E) Frequencies of Tregs among CD45^+^ cells or CD4^+^ T cells in 6421c2-bearing *Pdgfra^CreERT^*^2^*^/+;^I-Ab^fl/fl^* or *Pdgfra^+/+^;I-Ab^fl/fl^*mice receiving tamoxifen administration. Data are pooled from two independent experiments. Data are shown as mean ± SEM; *p<0.05; ****p<0.0001. Statistics were performed using the unpaired Student’s t-test.

## Notes

### Competing Interest Statement

The authors have declared no competing interest.

## References

1. Siegel, R. L., Kratzer, T. B., Wagle, N. S., Sung, H. & Jemal, A. Cancer statistics, 2026. CA Cancer J. Clin. 76, e70043 (2026).

2. Han, X. et al. Cancer-associated fibroblasts maintain critical pancreatic cancer cell lipid homeostasis in the tumor microenvironment. Cell Rep. 43, 114972 (2024).

3. Mucciolo, G. et al. EGFR-activated myofibroblasts promote metastasis of pancreatic cancer. Cancer Cell 42, 101–118.e11 (2024).

4. Liu, T., Zhou, L., Li, D., Andl, T. & Zhang, Y. Cancer-associated fibroblasts build and secure the tumor microenvironment. Front. Cell Dev. Biol. 7, 60 (2019).

5. Zhu, Z. et al. Tumour-reprogrammed stromal BCAT1 fuels branched-chain ketoacid dependency in stromal-rich PDAC tumours. Nat. Metab. 2, 775–792 (2020).

6. Ogier, C. et al. ANO6-mediated trogocytosis of cancer-associated fibroblasts promotes pancreatic cancer cell growth and immune suppression. *Nat*. Cancer 1– 22 (2026).

7. Kong, L. et al. Multimodal transcriptomics reveal myCAF-derived collagen signaling associated with impaired NK-cell cytotoxicity and immune exclusion in pancreatic cancer. J. Transl. Med. 24, (2026).

8. Öhlund, D. et al. Distinct populations of inflammatory fibroblasts and myofibroblasts in pancreatic cancer. J. Exp. Med. 214, 579–596 (2017).

9. Biffi, G. et al. IL1-induced JAK/STAT signaling is antagonized by TGFβ to shape CAF heterogeneity in pancreatic ductal adenocarcinoma. Cancer Discov. 9, 282– 301 (2019).

10. Elyada, E. et al. Cross-species single-cell analysis of pancreatic ductal adenocarcinoma reveals antigen-presenting cancer-associated fibroblasts. Cancer Discov. 9, 1102–1123 (2019).

11. Wang, Y. et al. Single-cell analysis of pancreatic ductal adenocarcinoma identifies a novel fibroblast subtype associated with poor prognosis but better immunotherapy response. Cell Discov. 7, 36 (2021).

12. Zhang, S. et al. Single cell transcriptomic analyses implicate an immunosuppressive tumor microenvironment in pancreatic cancer liver metastasis. Nat. Commun. 14, 5123 (2023).

13. Loveless, I. M. et al. Human pancreatic cancer single-cell atlas reveals association of CXCL10+ fibroblasts and basal subtype tumor cells. Clin. Cancer Res. 31, 756– 772 (2025).

14. Chen, X. et al. Single-cell resolution spatial analysis of antigen-presenting cancer-associated fibroblast niches. Cancer Cell 43, 2224–2240.e9 (2025).

15. Chen, K. et al. Complement-secreting CAFs are associated with better prognosis in pancreatic cancer: single-cell multiomics. Gut 75, 1367–1382 (2026).

16. Shi, Y. et al. Targeting LIF-mediated paracrine interaction for pancreatic cancer therapy and monitoring. Nature 569, 131–135 (2019).

17. Varveri, A. et al. Immunological synapse formation between T regulatory cells and cancer-associated fibroblasts promotes tumour development. Nat. Commun. 15, 4988 (2024).

18. Yagi, T. et al. Cancer-associated fibroblasts promote pro-tumor functions of neutrophils in pancreatic cancer via IL-8: potential suppression by pirfenidone. Cancer Immunol. Immunother. 74, 96 (2025).

19. Ringham, O. R. et al. A novel CAF population coordinates hyper-suppressive regulatory T cell recruitment and localization in lung cancer. Nat. Immunol. 27, 1899–1912 (2026).

20. Hiraoka, N., Onozato, K., Kosuge, T. & Hirohashi, S. Prevalence of FOXP3+ regulatory T cells increases during the progression of pancreatic ductal adenocarcinoma and its premalignant lesions. Clin. Cancer Res. 12, 5423–5434 (2006).

21. Tang, Y. et al. An increased abundance of tumor-infiltrating regulatory T cells is correlated with the progression and prognosis of pancreatic ductal adenocarcinoma. PLoS One 9, e91551 (2014).

22. Saleh, R. & Elkord, E. FoxP3+ T regulatory cells in cancer: Prognostic biomarkers and therapeutic targets. Cancer Lett. 490, 174–185 (2020).

23. Debesset, A. et al. TNFR2 blockade promotes antitumoral immune response in PDAC by targeting activated Treg and reducing T cell exhaustion. J. Immunother. Cancer 12, e008898 (2024).

24. Huang, H. et al. Mesothelial cell-derived antigen-presenting cancer-associated fibroblasts induce expansion of regulatory T cells in pancreatic cancer. Cancer Cell 40, 656–673.e7 (2022).

25. Maru, S. Y. et al. Antigen-presenting cancer-associated fibroblasts in murine pancreatic tumors differentially regulate T-cell phenotype and function. J. Immunol. 215, vkag165 (2026).

26. Cao, Z. et al. Interferon-γ-stimulated antigen-presenting cancer-associated fibroblasts hinder neoadjuvant chemoimmunotherapy efficacy in lung cancer. Cell Rep. Med. 6, 102017 (2025).

27. Kerdidani, D. et al. Lung tumor MHCII immunity depends on in situ antigen presentation by fibroblasts. J. Exp. Med. 219, e20210815 (2022).

28. Song, J. et al. Antigen-presenting cancer associated fibroblasts enhance antitumor immunity and predict immunotherapy response. Nat. Commun. 16, 2175 (2025).

29. Li, J. et al. Tumor cell-intrinsic factors underlie heterogeneity of immune cell infiltration and response to immunotherapy. Immunity 49, 178–193.e7 (2018).

30. Hingorani, S. R. et al. Trp53R172H and KrasG12D cooperate to promote chromosomal instability and widely metastatic pancreatic ductal adenocarcinoma in mice. Cancer Cell 7, 469–483 (2005).

31. Zhou, L. et al. Cancer-associated fibroblasts: origins, functional heterogeneity and therapeutic potential. Signal Transduct. Target. Ther. 11, 476 (2026).

32. Zhang, T., Ren, Y., Yang, P., Wang, J. & Zhou, H. Cancer-associated fibroblasts in pancreatic ductal adenocarcinoma. Cell Death Dis. 13, 897 (2022).

33. Principe, D. R. et al. TGFβ signaling in the pancreatic tumor microenvironment promotes fibrosis and immune evasion to facilitate tumorigenesis. Cancer Res. 76, 2525–2539 (2016).

34. Bockman, A. et al. Intratumoral regulatory T-cell ablation mediates tumor control systemically without autoimmunity. Cancer Res. 86, 4028–4042 (2026).

35. Zhang, C., et al. Intratumoral Treg cell ablation elicits NK cell-mediated control of CD8 T cell-resistant tumors. Sci. Immunol. 11, eadx4411 (2026).

36. Teng, M. W. L. et al. Conditional regulatory T-cell depletion releases adaptive immunity preventing carcinogenesis and suppressing established tumor growth. Cancer Res. 70, 7800–7809 (2010).

37. Joshi, N. S. et al. Regulatory T cells in tumor-associated tertiary lymphoid structures suppress anti-tumor T cell responses. Immunity 43, 579–590 (2015).

38. Plitas, G. et al. Regulatory T cells exhibit distinct features in human breast cancer. Immunity 45, 1122–1134 (2016).

39. Zhang, Y. et al. Regulatory T-cell depletion alters the tumor microenvironment and accelerates pancreatic carcinogenesis. Cancer Discov. 10, 422–439 (2020).

40. Weaver, J. D. et al. Differential expression of CCR8 in tumors versus normal tissue allows specific depletion of tumor-infiltrating T regulatory cells by GS-1811, a novel Fc-optimized anti-CCR8 antibody. Oncoimmunology 11, 2141007 (2022).

41. Van Damme, H. et al. Therapeutic depletion of CCR8+ tumor-infiltrating regulatory T cells elicits antitumor immunity and synergizes with anti-PD-1 therapy. J. Immunother. Cancer 9, e001749 (2021).

42. Villarreal, D. O. et al. Targeting CCR8 induces protective antitumor immunity and enhances vaccine-induced responses in colon cancer. Cancer Res. 78, 5340–5348 (2018).

43. Löhr, M. et al. Transforming growth factor-beta1 induces desmoplasia in an experimental model of human pancreatic carcinoma. Cancer Res. 61, 550–555 (2001).

44. Marshall, N. B. et al. NKG2C/E marks the unique cytotoxic CD4 T cell subset, ThCTL, generated by influenza infection. J. Immunol. 198, 1142–1155 (2017).

45. Tang, X. X., Shimada, H. & Ikegaki, N. Clinical relevance of CD4 cytotoxic T cells in high-risk neuroblastoma. Front. Immunol. 12, 650427 (2021).

46. Bravo, J. I., Kim, M. & Benayoun, B. A. Protocol for isolation of adult mouse ear pinnae-derived primary fibroblasts. STAR Protoc. 2, 100406 (2021).

47. Ivashkiv, L. B. IFNγ: signalling, epigenetics and roles in immunity, metabolism, disease and cancer immunotherapy. Nat. Rev. Immunol. 18, 545–558 (2018).

48. Zheng, Y. et al. Cancer-associated fibroblast signature can predict prognosis and therapeutic responses in skin cutaneous melanoma. Pigment Cell Melanoma Res. 38, e70005 (2025).

49. Ren, Y. et al. A multi-omics atlas of CAF subtypes reveals apCAF-M2 macrophage interactions driving immune resistance in glioma. PLoS One 20, e0329801 (2025).

50. Dauer, P. et al. Inactivation of cancer-associated-fibroblasts disrupts oncogenic signaling in pancreatic cancer cells and promotes its regression. Cancer Res. 78, 1321–1333 (2018).

51. Han, X. et al. Reversal of pancreatic desmoplasia by re-educating stellate cells with a tumour microenvironment-activated nanosystem. Nat. Commun. 9, 3390 (2018).

52. Krishnamurty, A. T. et al. LRRC15+ myofibroblasts dictate the stromal setpoint to suppress tumour immunity. Nature 611, 148–154 (2022).

53. Hutton, A. J. et al. Human lung fibroblasts present bacterial antigens to autologous lung Th cells. J. Immunol. 198, 110–118 (2017).

54. Koyama, M. et al. MHC class II antigen presentation by the intestinal epithelium initiates graft-versus-host disease and is influenced by the Microbiota. Immunity 51, 885–898.e7 (2019).

55. De Riva, A., Bourgeois, C., Kassiotis, G. & Stockinger, B. Noncognate interaction with MHC class II molecules is essential for maintenance of T cell metabolism to establish optimal memory CD4 T cell function. J. Immunol. 178, 5488–5495 (2007).

56. Kureshi, C. T. S. et al. Dendritic cell redundancy enables priming of anti-tumor CD4+ T cells in pancreatic cancer. Cancer Cell 44, 1381–1400.e9 (2026).

57. Kruse, B. et al. CD4+ T cell-induced inflammatory cell death controls immune-evasive tumours. Nature 618, 1033–1040 (2023).

58. Liu, X. The paradoxical role of IFN-γ in cancer: Balancing immune activation and immune evasion. Pathol. Res. Pract. 272, 156046 (2025).

59. Wang, S. et al. IFN-γ-driven UBE2D3 upregulation impairs antigen presentation pathways and anti-tumor immunity in pancreatic cancer. Nat. Commun. 16, 10733 (2025).

60. Kim, J. M., Rasmussen, J. P. & Rudensky, A. Y. Regulatory T cells prevent catastrophic autoimmunity throughout the lifespan of mice. Nat. Immunol. 8, 191– 197 (2007).

